# Chromatin-associated lncRNA-splicing factor condensates regulate hypoxia responsive RNA processing of genes pre-positioned near nuclear speckles

**DOI:** 10.1101/2024.10.31.621310

**Authors:** You Jin Song, Min Kyung Shinn, Sushant Bangru, Yu Wang, Qinyu Sun, Qinyu Hao, Pankaj Chaturvedi, Susan M. Freier, Pablo Perez-Pinera, Erik R. Nelson, Andrew S. Belmont, Mitchell Guttman, Supriya G. Prasanth, Auinash Kalsotra, Rohit V. Pappu, Kannanganattu V. Prasanth

## Abstract

Hypoxia-induced alternative splicing (AS) regulates tumor progression and metastasis. Little is known about how such AS is controlled and whether higher-order genome and nuclear domain (ND) organizations dictate these processes. We observe that hypoxia-responsive alternatively spliced genes position near nuclear speckle (NS), the ND that enhances splicing efficiency. NS-resident MALAT1 long noncoding RNA, induced in response to hypoxia, regulates hypoxia-responsive AS. MALAT1 achieves this by organizing the SR-family of splicing factor, SRSF1, near NS and regulating the binding of SRSF1 to pre-mRNAs. Mechanistically, MALAT1 enhances the recruitment of SRSF1 to elongating RNA polymerase II (pol II) by promoting the formation of phase-separated condensates of SRSF1, which are preferentially recognized by pol II. During hypoxia, MALAT1 regulates spatially organized AS by establishing a threshold SRSF1 concentration near NSs, potentially by forming condensates, critical for pol II-mediated recruitment of SRSF1 to pre-mRNAs.

## Introduction

The tumor microenvironment is under constant flux of oxygen, glucose, and other nutrients in a heterogeneous manner. Hypoxia, a state of low oxygen concentration, is a cellular stress that is frequently observed in solid tumors due to the uncontrolled proliferation of cancer cells and defective tumor vascularization^1^. Adaptation of cancer cells to hypoxia promotes tumor cell heterogeneity, cancer progression, and metastasis^2-4^. Cells respond to hypoxia by stabilizing the HIF (Hypoxia-inducible factor) family of transcription factors (TFs), leading to the increased expression of genes, controlling angiogenesis, glycolysis, and cancer cell invasion^4^. Besides transcription, hypoxia also activates differential gene expression by regulating co- and post-transcriptional processes, including alternative pre-mRNA splicing (AS)^5-9^. Hypoxia-responsive AS controls key processes such as oncogene activity, metabolic reprogramming, metastasis, and angiogenesis^7,9,10^. For example, hypoxia-induced AS of *VEGF-A* (vascular endothelial growth factor-A), the glycoprotein that promotes micro-vascularization and angiogenesis, generates pro-angiogenic VEGF-A isoforms^11,12^. At present, little is understood about the regulatory circuits controlling hypoxia-responsive AS.

The 4D nuclear organization (4 dimensions, including time and space) allows differential gene expression in response to various cues, such as cellular stress^13-15^. Such higher-order nuclear organization contributes to selective interactions among nucleic acids (DNA and RNA) and proteins, influencing protein modifications and localizations of proteins and RNAs in specific nuclear hubs, thereby contributing to differential gene expression. A feature of higher-order nuclear organization in mammalian cells is the presence of membrane-less nuclear domains (NDs), which are enriched with a unique subset of proteins and RNA. In general, NDs are formed via phase separation, a segregative phase transition driven by multivalent interactions between molecules to form distinct phases to create separate environments^16^. Nuclear speckles (NS) or speckles are one such NDs that form via phase separation and are enriched with RNA binding proteins (RBPs) and RNAs controlling pre-mRNA processing and mRNA export^17-20^. Studies indicate that NSs function as active hubs, enhancing transcription, pre-mRNA splicing efficiency, and mRNA export^17-20^. At present, it is not clear whether the 4D nuclear organization, and NDs, such as NSs, dictate hypoxia-responsive co- and post-transcriptional gene expression.

In the present study, we observed that the hypoxia-responsive AS is regulated by the higher-order nuclear organization and the nuclear speckle-enriched chromatin-associated long noncoding RNA, MALAT1 (Metastasis-associated lung adenocarcinoma transcript 1). The hypoxia-responsive genes are pre-positioned near NSs. MALAT1, induced in response to hypoxia, regulates hypoxia-responsive AS, mediated by SR-family of splicing factor 1 (SRSF1). Mechanistically, the binding of MALAT1 enhances the driving forces for the formation of SRSF1 phase-separated condensates to increase the concentration of SRSF condensates near hypoxia-responsive genes proximal to speckles. Furthermore, carboxy-terminal domain (CTD) of RNA pol II preferentially recognizes MALAT1-initiated SRSF1 condensates over SRSF1 free pool. Finally, MALAT1-promoted SRSF1 condensates are transferred to CTD to generate RNA pol II CTD-associated splicing condensates. In summary, during hypoxia, MALAT1 regulates AS of genes near NS by establishing the threshold concentration of SRSF1 condensates, required for RNA pol II-dependent recruitment of SRSF1 to pre-mRNAs.

## Results

### Hypoxia induces differential gene expression and alternative splicing of pre-mRNA

To understand the mechanisms controlling hypoxia-induced differential gene expression and alternative splicing (AS), we performed paired-end deep RNA-sequencing (>100 million paired reads/sample in duplicates) using poly A+ RNA from MCF7 (luminal sub-type of breast cancer) and HEK-293T (SV40 large T-antigen transformed human embryonic kidney) (Figure 1A) undergoing acute hypoxia (0.2% oxygen for 24 hrs) (please see STAR method). Principal component analysis confirmed that the data set from biological replicates was highly consistent in our RNA-seq data sets. We defined differentially expressed genes (DEGs) as genes that displayed absolute value (fold change) > 2.0 and FDR < 0.05, in statistical analysis. We observed elevated expression of HIF-target genes upon hypoxia in both cell lines (Figure 1B and Supplementary Figure 1Aa-b). Compared to the genome-wide distribution of the weighted gene expression changes, HIF1- and HIF2-induced genes showed significantly higher expression changes in response to hypoxia, thus indicating that HIF1 and HIF2 activate hypoxia-responsive genes in both cell lines (Supplementary Figure 1Aa-b). Both MCF7 and HEK293T showed dramatic, and cell type-specific differential expression of genes (DEG) in response to hypoxia (Figure 1C and Supplementary Figure 1B). Specifically, MCF7 and HEK293T showed differential expression of ∼3000 and 4000 genes respectively (Figure 1C, Supplementary Figure 1B) (Supplementary Tables 1-2). For example, hypoxic MCF7 and HEK293T cells showed elevated mRNA levels of 2138 and 2237 genes respectively. The hypoxia-responsive genes, for example, in MCF7 were enriched for several critical cancer-relevant pathways such as response to oxygen levels, stress, wounding, metabolism, cell signaling, and cell migration (Supplementary Figure 1C-D). Further, hypoxia-induced genes were associated with various types of cancer (Supplementary Figure 1E). Also, ∼30% hypoxia-responsive genes in MCF7 showed aberrant expression in tumor tissue samples from luminal breast cancer patients, as analyzed using the TCGA BrCa patient data set (Supplementary Figure 1F) (Supplementary Table 3) Next, we performed differential splicing analyses using rMATS with the RNA-seq datasets (see STAR methods for details). We defined hypoxia-responsive AS for splicing events with >= 0.15 IncLevelDifference (delta PSI) and P-value < 0.05 between normoxia and hypoxia samples. By this, we observed hypoxia-responsive AS of thousands of pre-mRNAs (Figure 1D and Supplementary Figure 1G) in both cell lines (Supplementary Tables 4-5). For example, hypoxic MCF7 cells showed ∼2100 hypoxia-specific AS splicing events in 1494 genes (1386 changes in cassette exons, 132 mutually exclusive exons, 251 retained-introns, 140 alternative 3’-splice sites, and 118 alternative 5’-splice sites) (Supplementary Tables 4-5). Alternatively spliced genes were involved in critical processes, such as regulation of stress response, RNA splicing, stress granule assembly, metabolic processes, and epithelial-mesenchymal transition (EMT) (Supplementary Figure 1H). A recent study reported that in HeLa cells hypoxia preferentially induced exon-skipping^21^. Like this report, HEK293T cells showed enhanced exon skipping in response to hypoxia (66% exon skipping; Supplementary Figure 1G). On the other hand, MCF7 showed only 36% hypoxia-responsive exon-skipping (Figure 1D), implying that hypoxia-responsive AS is regulated in cell type-specific manner. Interestingly, in MCF7, we observed minimal overlap between hypoxia-responsive differentially expressed and alternatively spliced genes (Figure 1E). This result implies that hypoxia-responsive AS is not a consequence of enhanced transcription, and that DEG and AS could be independently regulated.

**Figure 1.**
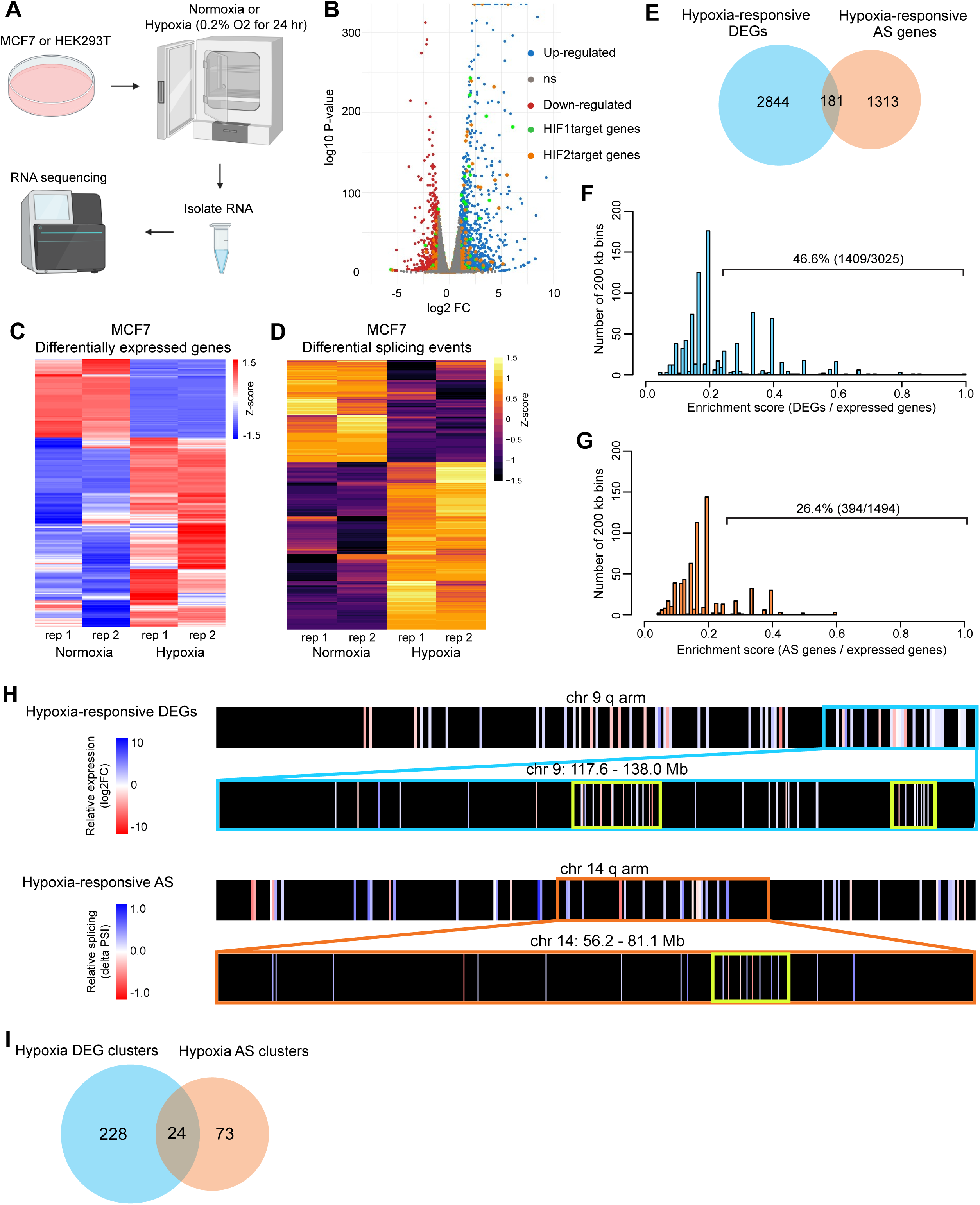
Hypoxia induces differential gene expression and alternative splicing of pre-mRNA in BrCa cells. (A) Schematic of hypoxia experimental set-up. (B) Volcano plot of differential gene expression during hypoxia in MCF7. Blue dots represent hypoxia up-regulated genes, red dots represent hypoxia down-regulated genes, green dots represent HIF1 target genes, orange dots represent HIF2 target genes, and gray dots represent the expressed genes. (C) Heatmap of hypoxia-responsive DEGs in MCF7. Genes (rows of heatmap) are hierarchically clustered using average-linkage clustering method. Color represents in Z-scores are computed per gene by subtracting the mean and then dividing by the standard deviation. Red and blue indicates up- and down-regulated genes respectively. (D) Heatmap of hypoxia-responsive AS events in MCF7. Splicing events (rows of heatmap) are hierarchically clustered using average-linkage clustering method. Color represents in Z-scores are computed per splicing event by subtracting the mean and then dividing by the standard deviation. Yellow indicates higher exon inclusion and violet indicates decreased exon inclusion. (E) Venn diagram showing overlap of hypoxia-responsive DEGs and AS genes. (F) Distribution of DEGs in 200 kb genomic bins with at least 5 annotated and expressed genes. Enrichment is calculated by dividing the number of hypoxia DEGs by the number of expressed annotated genes/genomic bin. Bins with enrichment score 0 not shown; full histogram in Supplementary figure 1I. (G) Distribution of AS genes in 200 kb genomic bins with at least 5 expressed annotated genes. Enrichment is calculated by dividing the number of hypoxia AS genes by the number of expressed annotated genes/ genomic bin. Bins with enrichment score 0 not shown; full histogram in Supplementary figure 1J. (H) Chromosome-wide heatmap of hypoxia-responsive DEG and AS genes. Blue and orange boxes highlight genomic regions containing clusters (highlighted in yellow) of differentially expressed and AS genes respectively. (I) Venn diagram showing the number of overlap of DEG and AS gene clusters. See also Supplementary Figure 1.

We next determined the genomic distribution of hypoxia responsive genes. We divided the MCF7 genome in 200 kb bins, calculated the enrichment of hypoxia-responsive genes over the annotated and expressed genes within such 200 kb genomic bins that contained five or more expressed genes (Figure 1F-G and Supplementary Figure 1I-J). The genome average for 200 kb bin is ∼1 annotated gene that was expressed in MCF7 cells. We defined a genomic bin as a hypoxia-responsive gene cluster if more than 25% of the genes within a bin are either differentially expressed or alternatively spliced upon hypoxia (Figure 1F-G; >0.25 in the X-axis). This cut-off was greater than the standard deviation of the genome average to identify gene clusters with significantly higher enrichment of hypoxia-responsive genes. As expected, most of the genomic bins did not contain or contained low numbers of hypoxia-responsive genes, and the bins that did contain hypoxia-responsive genes showed varying enrichment scores (mostly enriched between 0.2-0.6 for differentially expressed genes (DEGs) and 0.2-0.4 for AS genes) (Supplementary Figure 1I-J) (Supplementary Tables 8-9). We observed that 46.6% of hypoxia-responsive DEGs and 26.4% of AS genes were enriched in distinct gene clusters (Figure 1F-G). By plotting chromosome-wide heatmaps (Figure 1H and Supplementary Figure 1K), we observed that the hypoxia-responsive genes were non-randomly distributed in the chromosome (Supplementary Figure 1K). The length of the hypoxia-responsive gene clusters ranged from 200 kb (single bin) to 2.8 mb (multiple contiguous bins), containing ∼2-30 hypoxia-induced DEGs or AS genes. Based on this, we found 252 and 97 hypoxia-responsive differentially expressed and AS genomic clusters, respectively (Figure 1I). Interestingly, there was minimal overlap observed between differentially expressed and AS gene clusters (Figure 1I), indicating that a sub-population of hypoxia-responsive DEG and AS genes form distinct clusters in specific chromosome regions.

### Hypoxia-responsive genes pre-position near nuclear speckles (NS)

The clustering of genes in the linear genome for co-regulation has been observed previously, such as in the case of globin genes^22^ and olfactory receptor genes^23^. The mechanism by which the genes and gene clusters are brought to proximity in the 3D nuclear space usually involve higher-order chromatin organization and/or their association with various nuclear domains (NDs)^24^. Nuclear speckle (NS) or speckle is one such ND that is associated with active genes^25,26^, and co-ordinates transcription and RNA processing^27,28^. We therefore evaluated whether NSs influence hypoxia-responsive expression and AS of genes organized into genomic clusters. To determine this, we performed SON TSA-DNA-seq in normoxic and hypoxic MCF7 cells^29^. SON is a protein marker of NSs and is localized in the core of NSs^30,31^. TSA-DNA-seq utilizes a diffusible free radical reaction, in which the labeling efficiency on the DNA is correlated with the distance from the ND being labeled^29,32^. In our case, NS-localized SON allowed us to calculate the cytological distance of the genomic regions to the nearest speckles^29,32^. The relative distance of the genomic region to the NS is represented by the TSA genomic percentile score, wherein genomic regions are assigned genomic percentile scores based on the enrichment over input. Thus, a gene with a TSA percentile of 100 (TSA 100) is the relatively closest to NSs whereas a gene with a TSA percentile of 1 (TSA 1) is located farthest from speckles (Figure 2A-B).

**Figure 2.**
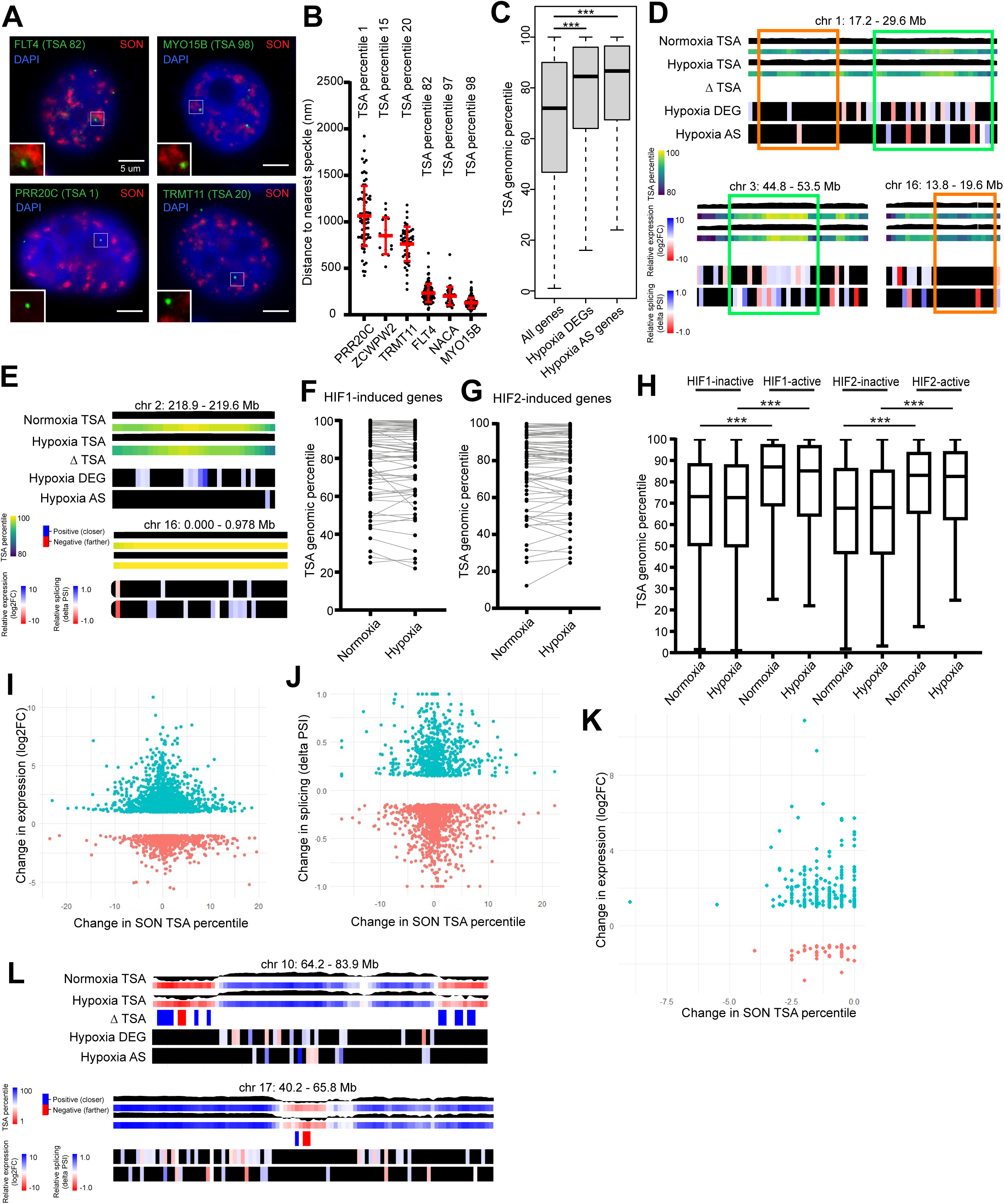
Hypoxia-responsive genes pre-position near nuclear speckle (NS) neighborhood in MCF7. (A) DNA FISH (green) and SON (red) immunostaining of speckle-near (high TSA percentile) and speckle-far (low TSA percentile) genes. DNA is counter-stained with DAPI (blue). (B) Scatter plot showing quantification of distance between DNA FISH foci to the nearest speckle for genes in different TSA percentiles. Error bars showing mean and standard deviation. (C) Box plot showing TSA percentiles of all expressed genes, hypoxia DEGs, and hypoxia AS genes. Box representing 25^th^ to 75^th^ percentile with line indicating mean value. Whiskers representing the min and max value. (*) P ≤ 0.05, (**) P ≤ 0.01, (***) P ≤ 0.001 by two-tailed student’s t-test. (D) Three representative chromosome regions enriched (green box) and depleted (orange) of hypoxia-responsive genes highlighted on the genome-wide heatmap of TSA percentile score. First and second row represents TSA percentile scores across the region in normoxia. Third and fourth rows represent TSA percentile scores across the region in hypoxia. First and third rows represent the SON TSA-seq mapping wherein a peak indicates NS proximal region and valley represents NS far region. Second and fourth rows show heatmaps representing the TSA genomic percentile wherein yellow indicates TSA genomic percentile 100 and violet indicates TSA percentile 80. The fifth row (ΔTSA) indicates regions of movement or significant change in genomic regions in TSA percentile from normoxia to hypoxia. Blue and red indicates genomic regions moving closer or farther from NSs during hypoxia. Sixth row indicates hypoxia DEGs, with blue indicating up-regulation and red indicating down-regulation. Seventh row indicates hypoxia AS genes, with blue and red indicating increased and decreased exon inclusion respectively. (E) Hypoxia DEG and AS clusters on the genome-wide heatmap of TSA percentile score. (F) Pairwise plot showing SON-TSA genome percentile of HIF1-induced genes in normoxia and hypoxia. (G) Pairwise plot showing SON-TSA genome percentile of HIF2-induced genes in normoxia and hypoxia. (H) Box plot showing SON-TSA genome percentiles of HIF1-bound transcriptionally-inactive and -active genes and HIF2-bound inactive and active genes. Box representing 25^th^ to 75^th^ percentile with line indicating mean value. Whiskers representing the min and max value. (*) P ≤ 0.05, (**) P ≤ 0.01, (***) P ≤ 0.001 by two-tailed student’s t-test. (I) Scatter plot showing change in TSA percentile correlated with change in expression. (J) Scatter plot showing change in TSA percentile correlated with change in splicing. (K) Scatter plot showing SPAD gene change in TSA percentile correlated with change in expression. (L) Hypoxia-responsive genes highlighted on the genome-wide heatmap of TSA percentile score. First and second rows represent TSA percentile scores across the region in normoxia. Third and fourth rows represent TSA percentile scores in hypoxia. First and third rows represent the SON TSA-seq mapping wherein a peak indicates NS proximal region and valley represents NS far region. Second and fourth rows show heatmaps representing the TSA genomic percentile wherein blue indicates high TSA percentile and red indicates low TSA percentile. The fifth row (ΔTSA) indicates regions of movement or significant change in TSA percentile from normoxia to hypoxia. Blue and red indicates regions moving closer and farther from speckles during hypoxia respectively. Sixth row indicates hypoxia DEGs, with blue and red indicating up-regulated and down-regulated genes respectively. Seventh row indicates hypoxia AS genes, with blue and red indicating increased and decreased exon inclusion respectively. See also Supplementary Figure 2.

By performing SON-TSA-seq, we determined the relative distribution of hypoxia-induced and AS genes in the MCF7 cell nucleus. We observed that hypoxia-responsive DEG as well as alternatively spliced (AS) genes were located significantly closer to NSs compared to hypoxia non-responsive, constitutively expressed and -spliced genes (Figure 2C & Supplementary Figure 2A). Hypoxia-responsive genes displayed a higher mean TSA-score of ∼85 compared to constitutively transcribed and -spliced genes (mean TSA-score of ∼70) (Figure 2C) (Supplementary Tables 10-11). DNA-FISH further confirmed SON TSA-seq data, showing close speckle proximity distribution of genes with high TSA score (*FLT4* and *MYO15B, NACA*) compared to the ones with low SON TSA-seq score (*PRR20C, TRMT11, ZCWPW2*) (Figures 2A-B). Further, chromosome-wide data representation also indicated that hypoxia-responsive genes enriched in speckle proximity (Figures 2D-E, L & Supplementary Figure 2A & F). Please note that in Figure 2D-E, the heatmap for the speckle-proximal regions are represented by TSA genomic percentile 100 in yellow and TSA genomic percentile 80 in violet to show better resolution (Figure 2D-E). In the rest of the chromosome-wide heatmaps (for example, Figure 2L) showing the full range of TSA genomic percentiles are represented by TSA genomic percentile 100 in blue and TSA genomic percentile 1 in red. Based on the genomic distribution of all the expressed genes compared to only the hypoxia-responsive genes, the hypoxia-responsive genes were not evenly distributed across the genome but showed a higher enrichment in the previously defined SPADs (speckle-associated domains with TSA score between 95-100)^29^ (Supplementary figure 2B-E). Also, within each speckle-proximal region (TSA score between 80-100), hypoxia-responsive genes formed sub-clusters (Figure 2D and Supplementary figure 2B-E). For example, in Figure 2D, the green rectangle within a speckle proximal region was enriched with hypoxia responsive genes. On the other hand, the orange rectangle within the same speckle proximal region contained low density of hypoxia responsive genes. This implies that besides gene organization around NS neighborhood, additional factors influence selective hypoxia-responsive transcription and AS of genes. Also, as pointed out earlier (Figure 1I) hypoxia-responsive DE (Figure 2Ea) and AS (Figure 2Eb) gene clusters were organized as distinct non-overlapping clusters around NSs.

Recent studies reported stress-induced repositioning of genes close to NSs for gene activation and differential splicing^15,33-35^. We therefore examined such repositioning of hypoxia-induced or -alternatively spliced genes to NS neighborhood. We first analyzed potential changes in gene positioning of HIF1- and HIF2-target genes, which were induced in the hypoxic MCF7 cells (Figure 1B). Most of the HIF1- and HIF2-induced genes remained positioned near NSs in normoxic and hypoxic cells as they displayed high TSA-score (Figure 2F-H and Supplementary Figure 2F). Twenty-four hrs of acute hypoxia did not show significant change in repositioning of most if not all the HIF-induced genes towards speckles (Figure 2F-H). DNA-FISH analyses further confirmed speckle proximity localization of HIF-target genes in both normoxic and hypoxic cells (Supplementary Figure 2G; please see the quantification of the DNA-FISH data in Figure 6A). Based on the published HIF ChIP-seq data, in hypoxic MCF7 cells, HIF1 and HIF2 are bound to the promoters of 503 and 661 genes^36^ (Supplementary Tables 6-7). Acute hypoxia (0.2% O_2_ for 24 hrs) in MCF7 cells showed significant upregulation of ∼115 of HIF-target genes. We determined whether the spatial positioning of these 115 genes in the 3D nuclear space, especially near NS, helps them to be hypoxia-responsive compared to the rest of the HIF target genes, which did not respond to acute hypoxia for 24 hrs. We observed that hypoxia-induced HIF target genes (HIF1- or 2-active) were pre-positioned closer to NSs, compared to the rest of the non-induced HIF target genes (HIF1- or 2-inactive) (Figure 2H). Most importantly, hypoxia-induced HIF-active genes were pre-positioned in NS proximity, even in normoxic cells (a time window where these genes were not induced) and did not show change in gene position during hypoxia (Figure 2H). These results indicate that in MCF7 cells, HIF-target genes that are pre-positioned at speckle proximity in normoxia cells respond more efficiently to hypoxia.

Next, we plotted the change in SON TSA-genomic percentile with changes in differential expression and AS of all the hypoxia responsive genes (Figure 2I-J). Based on these analyses, we did not observe any positive correlation between gene movement towards NSs and hypoxia-responsive gene activation and AS and in MCF7 cells that underwent acute hypoxia for 24 hrs (Figure 2I-J). We then examined whether the genes that indeed displayed a significant change in hypoxia-responsive repositioning, (defined based on the significant change in TSA genomic percentile [difference greater than variation between biological replicates and P-value <= 0.05]), showed enhanced expression or AS (including, exon-inclusion or -exclusion) (Supplementary Figure 2H-K). Based on this criterion, we observed that out of the 2138 hypoxia-induced genes in MCF7, only 76 genes (∼3.5%) showed significant change in gene position. Similarly, only ∼1.9% of the hypoxia-responsive AS genes displayed gene repositioning. Most importantly, hypoxia-responsive gene movement towards or away from NSs did not correlate with their induced expression or AS (Supplementary Figure 2H-K).

By comparing the expression of genes within the SPADs (95-100 TSA genomic percentile) in various cell lines, an earlier study reported positive correlation between gene expression and their localization at NS proximity^32^. We determined whether there exists any positive correlation between SPAD-associated gene movement (P-value <= 0.05) towards NSs and hypoxic-responsive induction (Figure 2K). Again, we did not observe any such positive correlation, rather the genes within the SPAD that were induced in response to hypoxia moved away from NSs (Figure 2K; 0 to -2.5 change in TSA score). In general, most of the genomic regions that displayed significant changes in position (indicated by ΔTSA) post-hypoxia were positioned in speckle-far regions (chromosome regions marked shades of red) and did not contain hypoxia-responsive genes (Figure 2L). In summary, we conclude that in MCF7 cells, the hypoxia-responsive expression and AS are not correlated with gene movement towards NSs. The hypoxia-responsive genes, organized in distinct genomic “clusters” are pre-positioned near NSs.

### MALAT1 regulates the hypoxia response in breast cancer cells

A significant part of the human genome is transcribed into noncoding RNAs (ncRNAs) that are retained in the nucleus, where they interact with chromatin, hence named caRNAs (<u>chromatin-associated</u> <u>RNAs</u>). CaRNAs regulate gene expression by modifying chromatin structure^37^. Hypoxia induces the expression of hundreds of caRNAs^38,39^. At present, we know the function of only a handful of caRNAs in hypoxia-responsive gene expression and RNA processing^40-42^. Specifically, we focused on the involvement of caRNAs in hypoxia-responsive AS. RNA-seq analyses revealed that hypoxia induced the differential expression of 921 and 1357 lncRNAs in MCF7 and HEK293T cells respectively. One hundred and thirty-eight of them were differentially expressed in both the cell lines (Figure 3A) (Supplementary Table 12). MALAT1 (Metastasis-associated lung adenocarcinoma transcript 1) was identified as one of the most abundant HIF-target chromatin-associated lncRNAs that was induced several fold in response to hypoxia in both MCF7 and HEK293T cells (Figure 3B-C; Supplementary Figure 3A-B)^43^. Hypoxic MCF7 cells contain ∼15,000 copies of MALAT1/nucleus (Figure 3B and Supplementary Figure 3B). Earlier studies, including studies from our laboratory have demonstrated the role of MALAT1 in tumor progression and metastasis^44-47^. Luminal sub-type breast cancer (BrCa) patients with high MALAT1 expression showed reduced overall survival (Supplementary Figure 3C). To determine the impact of hypoxia-responsive induction of MALAT1 in BrCa progression, we examined various hypoxia-induced phenotypes in control, MALAT1-depleted (using modified antisense oligonucleotides) or MALAT1 knockout (KO) (using CRISPR-based approach) cells (Supplementary Figure 3D)^46,48-50^. Hypoxia enhances cancer cell migration, invasion, metastasis and EMT^51^. We observed that MALAT1 promoted the hypoxia-responsive BrCa cell migration and invasion (Figure 3D and 3E). MALAT1 KO MDA-MB-231 cells also showed reduced hypoxia-responsive wound-healing capacity (Figure 3F). Further, MALAT1-depleted non-tumorigenic mammary epithelial MCF10a cells failed to activate hypoxia-responsive wound-induced EMT, as observed by reduced vimentin-positive cells at the edge of a wound upon MALAT1 depletion (Supplementary Figure 3E). Hypoxic cells undergo metabolic reprogramming due to the decreased oxygen levels by relying on anaerobic glycolysis, thus synthesizing lactic acid as the product^52^. MALAT1-depleted cells showed defects in hypoxia-induced lactic acid secretion, thus implying its involvement in metabolic reprogramming (Figure 3G). We found that MALAT1 KO cells transplanted into the fat pad of nude mice (n =10) displayed decreased tumor progression and tumor angiogenesis (Figure 3H-I & Supplementary Figure 3F). Finally, MALAT1 KO hypoxic BrCa cells, compared to WT cells failed to efficiently colonize the lungs (metastasize) after a tail-vein graft (Supplementary Figure 3G). Results from these phenotypic assays imply that MALAT1 plays an important role in the hypoxic response of BrCa cells.

**Figure 3.**
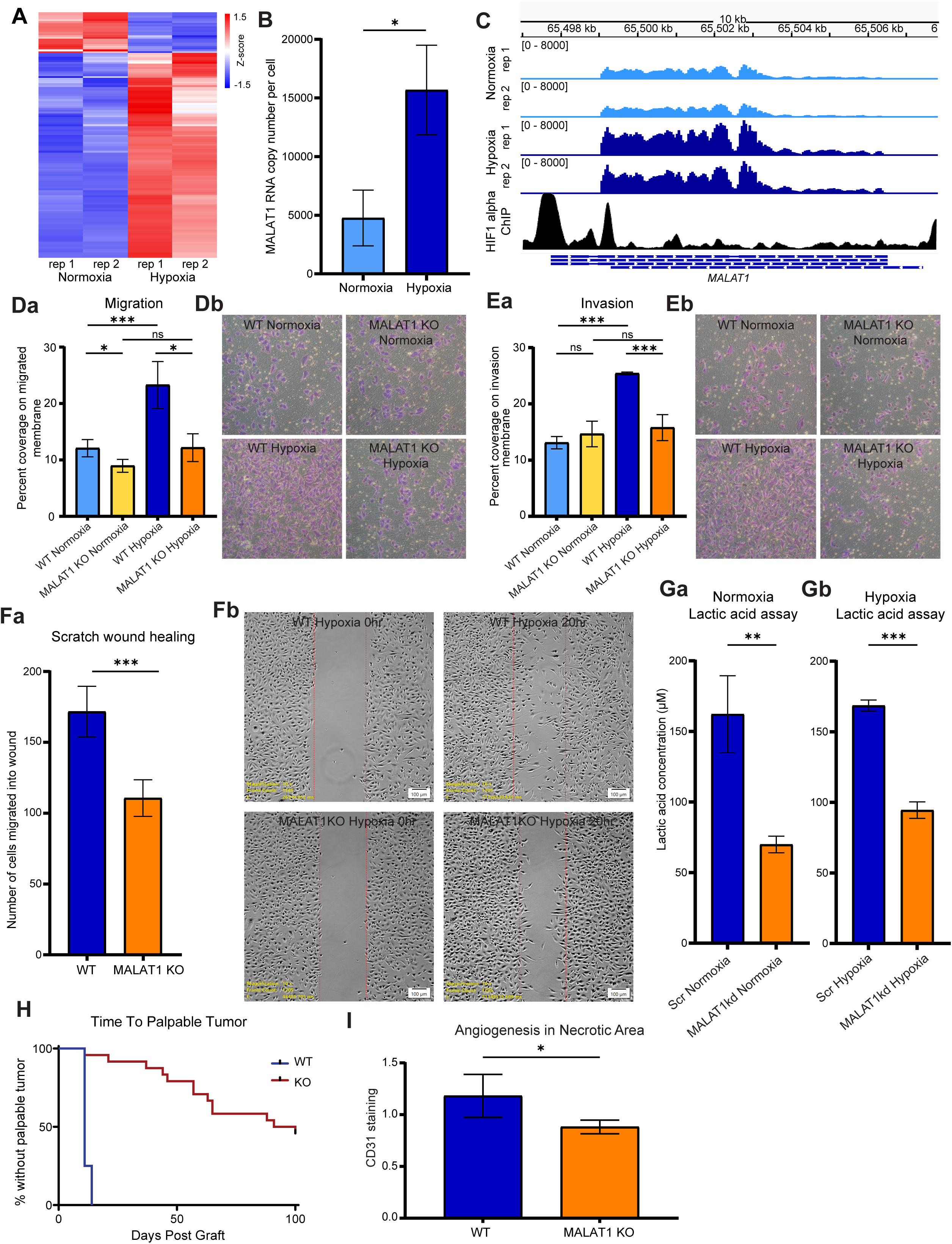
MALAT1, induced during hypoxia, regulates hypoxia-responsive changes in cancer cells. (A) Heatmap of hypoxia-responsive lncRNAs. Red and blue indicates and up- and down-regulated lncRNAs in MCF7. (B) Transcript copy number analysis of MALAT1 in normoxic and hypoxic MCF7 cells. (C) IGV representation of MALAT1 locus in normoxic and hypoxic MCF7 cells. HIF1-alpha ChIP data shown from hypoxic MCF7 cells (Da-b) Transwell migration assay of MDA-MB-231 wildtype and MALAT1 KO cells in normoxia and hypoxia. (E) Transwell invasion assay of MDA-MB-231 wildtype and MALAT1 KO cells in normoxia and hypoxia. (F) Live scratch wound healing assay of MDA-MB-231 wildtype and MALAT1 KO cells in hypoxia. (G) Lactic acid secretion assay of scramble control (Scr) and MALAT1kd MCF7 cells in normoxia and hypoxia. (H) Primary orthotopic tumor growth of wildtype and MALAT1 KO MDA-MB-231 mouse xenograft. Data is plotted as time from graft to first detection of a palpable tumor (N=10; no mice were censored). (I) Tumor angiogenesis measured by CD31 staining from wildtype and MALAT1 KO HEK293T tumors. Error bars (B, Da, Ea, Fa Ga-b, and I) showing standard deviation from three biological replicates. (*) P ≤ 0.05, (**) P ≤ 0.01, (***) P ≤ 0.001 by two-tailed student’s t-test. See also Supplementary Figure 3.

### MALAT1 regulates the differential expression and AS of hypoxia-responsive genes

In the nucleus, MALAT1 is enriched at the outer shell of the NS, where the hypoxia-responsive genes are distributed^31,50,53^. MALAT1 is known to interact with transcriptionally active and alternatively spliced genes^54,55^. We hypothesize that hypoxia-induced MALAT1 regulates the differential expression and AS of hypoxia-responsive genes near NSs. We determined changes in hypoxia-responsive expression and AS in control and MALAT1-depleted (transient by ASOs) or MALAT1 KO (by CRISPR) cells (both in MCF7 and HEK293T) by performing polyA+ RNA-seq RNA-seq following differential expression and splicing analyses (Supplementary Tables 13-17). Specifically, differential expression or AS events observed in the control, but not in the MALAT1-depleted hypoxic cells were quantified as MALAT1-dependent events. We found that ∼70% and 55% of the hypoxia-responsive gene expression changes were deregulated in MALAT1-depleted MCF7 and HEK293T cells, respectively (Figure 4A-B and Supplementary Figure 4A-B). MALAT1-dependent hypoxia-responsive genes regulated pathways relevant to hypoxia, such as response to hypoxia and wounding, angiogenesis and EMT (Supplementary Figure 4C).

**Figure 4.**
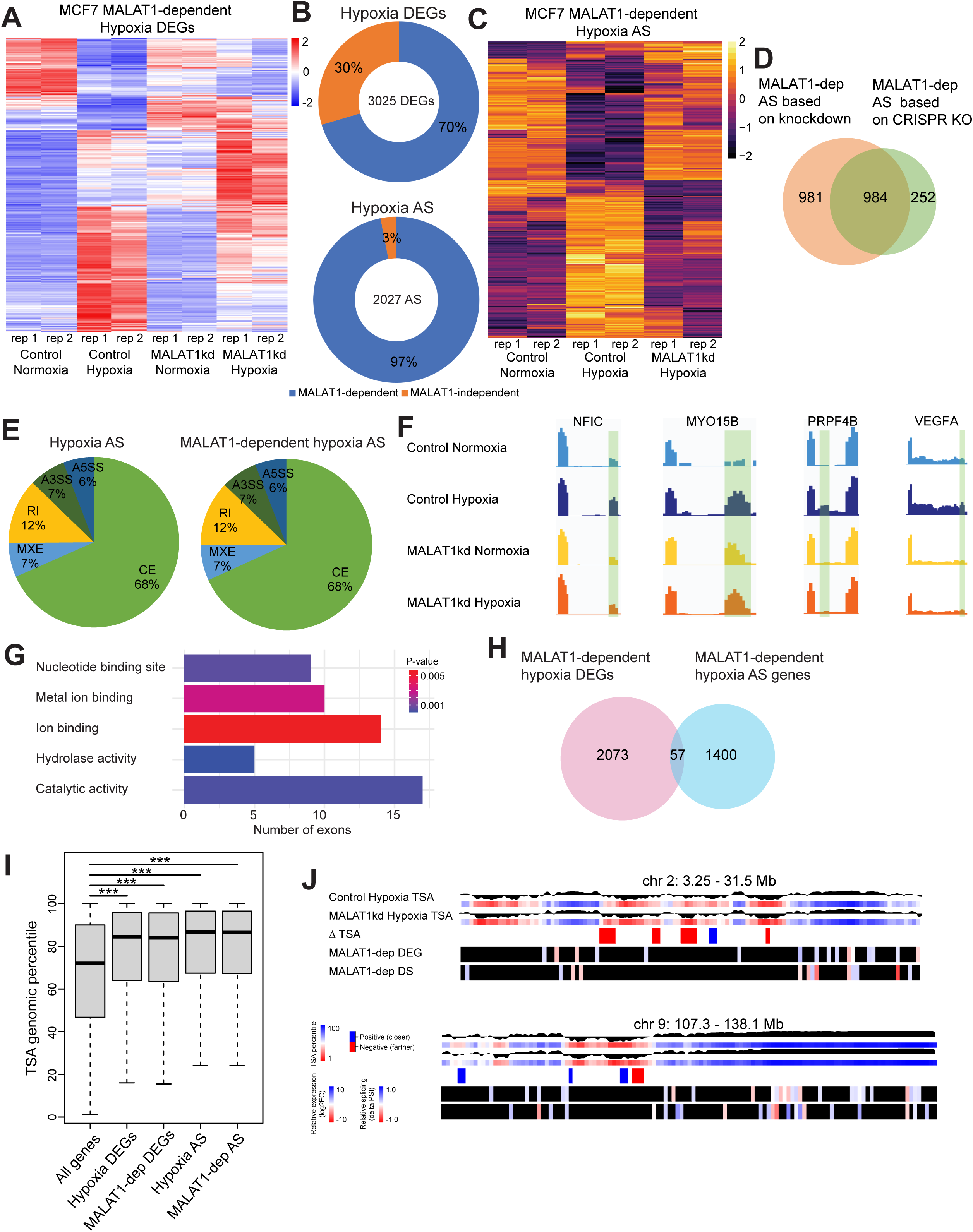
MALAT1 regulates hypoxia-responsive differential expression and AS in MCF7. (A) Heatmap of hypoxia-responsive DEGs in control and MALAT1kd MCF7 cells. Genes (rows of heatmap) are hierarchically clustered using average-linkage clustering method. Color represented in Z-scores are computed per gene by subtracting the mean and then dividing by the standard deviation. Red and blue indicate up- and down-regulated genes respectively. (B) Pie charts indicating percentage of hypoxia-responsive DEGs and AS events dependent on MALAT1. (C) Heatmap of hypoxia-responsive AS events in control and MALAT1kd MCF7 cells. Splicing events (rows of heatmap) are hierarchically clustered using average-linkage clustering method. Color represented in Z-scores are computed per splicing event by subtracting the mean and then dividing by the standard deviation. Yellow and violet indicate exon-inclusion and exclusion respectively. (D) Venn diagram showing the overlap of the MALAT1-dependent AS events from MALAT1 kd and MALAT1 KO MCF7. (E) Pie chart indicating classifications of the modes of alternative splicing. (F) Examples of MALAT1-dependent AS events. The highlighted exons show changes in MALAT1-depleted hypoxic cells. (G) Top five exon annotations of the MALAT1-regulated hypoxia AS exons. (H) Venn diagrams showing overlap of the MALAT1-dependent DEGs and AS genes. (I) Box plot showing TSA genome percentile of hypoxia-responsive and MALAT1-dependent genes. Box representing 25^th^ to 75^th^ percentile with line indicating mean value. Whiskers representing the min and max value. (*) P ≤ 0.05, (**) P ≤ 0.01, (***) P ≤ 0.001 by two-tailed student’s t-test. (J) Genome-wide distribution of MALAT1-dependent DEGs and AS genes from a chromosome region. First and second rows represent TSA percentile scores across the region in control hypoxia. Third and fourth rows represent TSA percentile scores across the region in MALAT1kd hypoxia. First and third rows represent the SON TSA-seq mapping wherein a peak indicates NS proximal region and valley represents NS far region. Second and fourth rows show heatmaps representing the TSA genomic percentile wherein blue and red indicate high and low TSA percentiles respectively. The fifth row indicates regions of significant change in TSA percentile (ΔTSA) between control hypoxia and MALAT1kd hypoxia. Blue and red indicate regions moving closer and farther from speckles in MALAT1kd respectively. Sixth row indicates MALAT1-dependent hypoxia DEGs, with blue and red indicating up- and down-regulated genes respectively. Seventh row indicates MALAT1-dependent hypoxia AS genes, with blue and red indicating increase and decreased exon inclusion respectively. See also Supplementary Figure 4.

Aside from gene expression, we also found that 97% and 82% of hypoxia-responsive AS events in MCF7 and HEK293T cells respectively were dysregulated upon MALAT1-knockdown (Figure 4B-C & Supplemental Figure 4B & D) (Supplementary Tables 14 & 16). Hypoxic MCF7 cells depleted for MALAT1, transiently (modified antisense DNA oligonucleotides) or permanently (CRISPR-KO), showed a significant overlap of concordant AS changes (Figure 4D) (Supplementary Tables 14 & 17). In general, MALAT1-depleted cells showed defects in all the annotated forms of AS (Figure 4E & Supplementary Figure 4E). MALAT1 influenced the hypoxia responsive AS of genes (such as *VEGF-A*, *NFIC*) (Figure 4F), controlling vital functions such as RNA metabolic processes and response to hypoxia (Supplementary Figure 4F). Further, exon annotation data analyses revealed that MALAT1-regulated exons were enriched for protein domains, containing nucleotide and ion binding and various enzymatic activities (Figure 4G). Interestingly, we observed minimal overlap between MALAT1-dependent differentially expressed and alternatively spliced genes in both cell lines (Figure 4H & Supplementary Figure 4G) (Supplementary Tables 13-16), potentially implying independent regulatory mechanisms. In summary, our results indicate that MALAT1 regulates hypoxia-responsive AS.

Next, we assessed whether MALAT1 influences the speckle proximal positioning of hypoxia-responsive genes. We determined the localization of MALAT1-dependent hypoxia-induced and AS genes in control and MALAT1-depleted hypoxic MCF7 cells by performing SON TSA-seq. MALAT1-dependent hypoxia-induced and -AS genes continued to pre-position near speckles in normoxic and hypoxic cells in presence or absence of MALAT1 (Figure 4I and Supplementary Figure 4I), implying that MALAT1 does not seem to influence the distribution of hypoxia-responsive genes. Both overall hypoxia-responsive genes and MALAT1-dependent hypoxia-responsive genes had significantly higher TSA genomic percentile scores compared to the genome average (Supplementary Figure 4I). Interestingly, MALAT1-depleted hypoxic cells did show re-positioning of several genes (compared to control cells), most of which located in speckle distal regions (Figure 4J and Supplementary Figure 4H; see ΔTSA). However, these genes did not show hypoxia-responsive DEG or AS (Figure 4J and Supplementary Figure 4H), indicating lack of correlation between MALAT1-dependent gene positioning and expression during hypoxia.

### MALAT1 enhances the specificity of SRSF1 binding to its target transcripts

We hypothesize that MALAT1 regulates hypoxia-responsive AS by modulating the activity of a subset of MALAT1-interacting and NS resident RBPs/splicing factors. Motif scanning analyses showed that >100s RBPs potentially interact with MALAT1 (Supplementary figure 5A; Supplementary Table 18). Earlier studies reported interaction of several dozens of RBPs, including splicing factors, with MALAT1^56-58^. We initially assessed whether the MALAT1-regulated AS exons and the nearby intron-exon junctions showed enrichment for MALAT1-interacting RBP/s. To do this, we calculated the enrichment of RBPs on the MALAT1-regulated exons and the constitutively-spliced (CS) exons (as negative control) in hypoxic cells (please see STAR method for details) using the publicly available RBP-binding datasets for 392 RBPs from the cisBP-RNA database (Supplementary Tables 19-20) (Figures 5A-B)^59^. Several splicing factors, including SR-family of splicing factors (SRSF1 and SRSF9), and SR-like proteins such has U1-70K, a component of U1 snRNP and an SRSF1 interactor, showed preferential enrichment on the MALAT1-regulated exons, compared to constitutively spliced exons (Figure 5A-B). We and others have previously shown that MALAT1 regulates the localization and activity of SRSF1^47,50,60,61^. SRSFs are known to regulate hypoxia-responsive AS, but the underlying regulatory mechanism is yet to be determined^7,10,21^. We therefore focused our efforts on SRSF1 and determined how MALAT1regulates SRSF1-mediated AS during hypoxia. Each MALAT1 RNA contained ∼100 SRSF1 binding motifs as observed by scanning motif analysis (Supplementary Figure 5B). Further, SRSF1 eCLIP-seq (<u>e</u>nhanced <u>c</u>rosslinking and immuno<u>p</u>recipitation-seq) in normoxic and hypoxic MCF7 and HEK293T cells revealed that SRSF1 extensively interacted with the entire length of MALAT1 RNA (Figure 5C and Supplementary Figure 5C). We detected ∼100 SRSF1 eCLIP cross-linked sites on MALAT1 RNA, implying that many SRSF1 proteins bind to each MALAT1 RNA. Furthermore, SRSF1-depleted hypoxic HEK293T cells showed changes in hypoxia-responsive AS (6570 AS events) (Figure 5D) (Supplementary Table 22). Interestingly nearly one-third (34%) of MALAT1-dependent hypoxia-responsive AS events (1821 out of 5370 AS events) showed concordant changes upon SRSF1 depletion (Figures 5D-F), thus indicating that a significant fraction of hypoxia-responsive AS events is controlled via MALAT1/SRSF1 axis.

**Figure 5.**
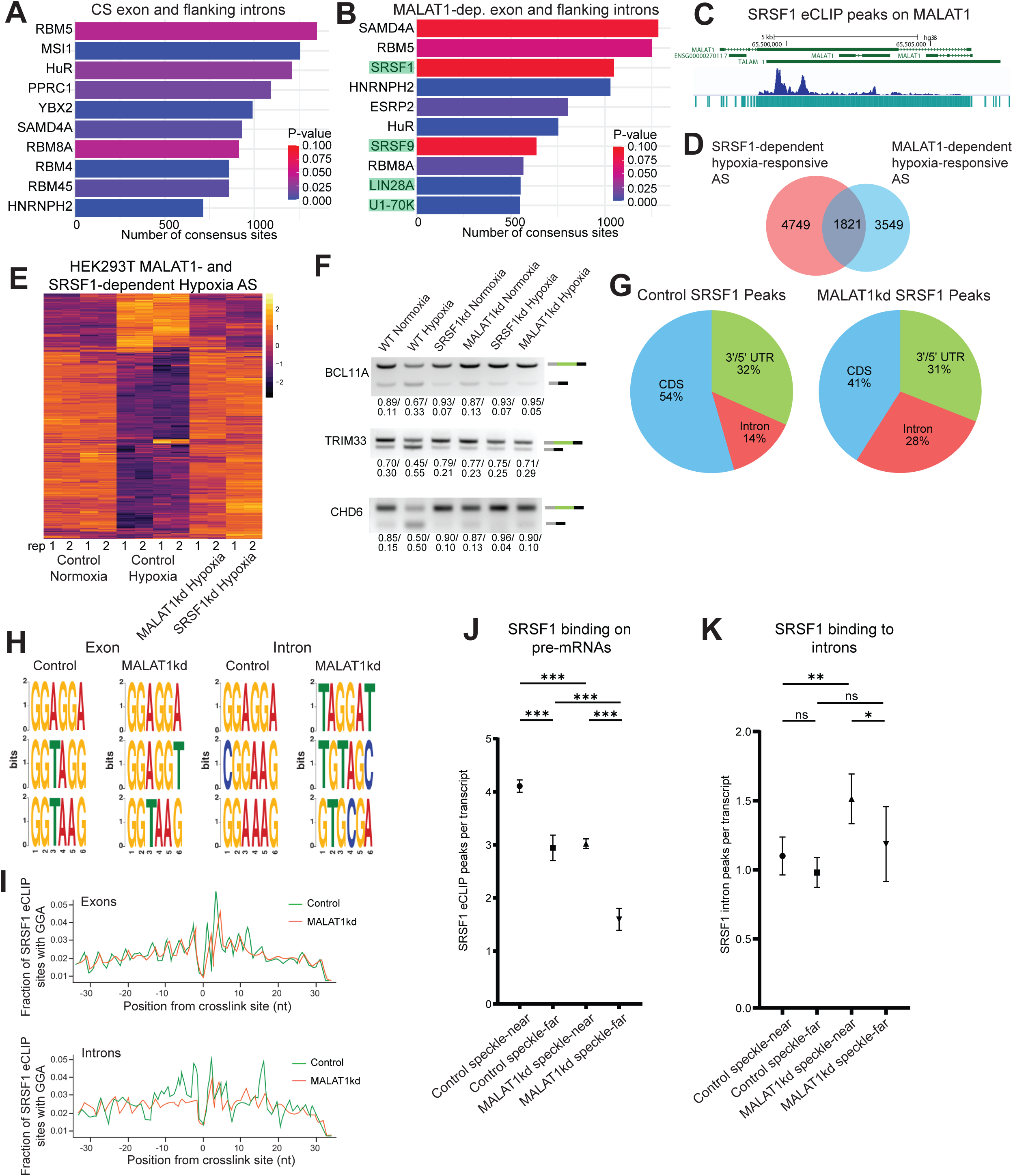
MALAT1 regulates the SRSF1 binding to its target transcripts. (A) Top ten RBP motif enrichment on constitutively spliced regions within hypoxic MCF7. Bar length indicates number of sites and color indicates P-value. (B) Top ten RBP motif enrichment on MALAT1-dependent spliced regions in hypoxic MCF7. Bar length indicates number of sites and color indicates P-value. Highlighted RBPs show preferential enrichment only in MALAT1-dependent spliced sequences (C) SRSF1 eCLIP peaks on MALAT1 locus. Middle plot showing normalized SRSF1 eCLIP alignment and bottom plot showing called peaks on *MALAT1* locus. (D) Venn diagram showing overlap of MALAT1- and SRSF1-dependent hypoxia AS events in HEK293T. (E) Heatmap of MALAT1- and SRSF1-dependent hypoxia AS events. Yellow indicates increased exon inclusion and violet indicates decreased exon inclusion. (F) RT-PCR of MALAT1- and SRSF1-dependent hypoxia AS events. (G) Pie chart showing SRSF1 eCLIP-seq peak annotation in control and MALAT1-depleted hypoxic MCF7. (H) Nucleotide enrichment analysis of SRSF1 binding sites from eCLIP-seq analysis. (I) Enrichment of GGA motif around SRSF1 crosslinking sites in control and MALAT1-depleted hypoxic MCF7 SRSF1 eCLIP-seq data. (J) SRSF1 eCLIP peak number per transcript in speckle-near and speckle-far genes in control and MALAT1-depleted MCF7. (K) SRSF1 eCLIP peak number in intronic regions per transcript in speckle-near and speckle-far genes in control and MALAT1-depleted samples. Data represented in J & K as mean +-SEM. (*) P ≤ 0.05, (**) P ≤ 0.01, (***) P ≤ 0.001 by two-tailed student’s t-test. See also Supplementary Figure 5.

To evaluate the mechanism by which MALAT1 regulates SRSF1-mediated AS, we determined transcriptome-wide SRSF1 binding to pre-mRNA in control and MALAT1-depleted normoxic and hypoxic cells by eCLIP-seq. SRSF1 eCLIP reads were processed to identify unique cross-link events^62^. SRSF1 eCLIP-seq revealed that SRSF1 bound to ∼30,000 and ∼20,000 sites 9655 & 5055 target genes in hypoxic MCF7 and HEK293T cells, respectively (Supplementary Tables 23-24). SRSF1 bound mostly to the exons, comprising both coding exons and UTR sequences^62-65^ (Figure 5G). Interestingly, MALAT1-depleted hypoxic cells altered SRSF1 binding preferences. We observed a small but significant decrease in the binding of SRSF1 to the coding exons in MALAT1-depleted hypoxic cells (54% in control versus 41% in MALAT1-depleted cells) (Figure 5G). Strikingly, MALAT1-depleted hypoxic cells showed a two-fold increase in SRSF1 binding to introns, compared to control cells (14% in control versus 28% in MALAT1-KD) (Figure 5G). Next, we performed a sequence enrichment analysis in the SRSF1 eCLIP-seq data sets to identify the SRSF1 binding sequence motifs in control and MALAT1-depleted cells. SRSF1 preferentially recognizes the consensus sequences of multiple GGA repeats^66^. As expected, GGA- and similar sequence repeats were enriched in the SRSF1 eCLIP-seq data within the exonic regions of control and MALAT1-depleted cells (Figure 5H). Similar GGA-repeat sequence enrichment was also observed in the SRSF1 cross-linked sites within the introns of control hypoxic cells (Figure 5H). However, upon MALAT1-depletion, SRSF1 failed to preferentially bind to the consensus GGA-repeats within the introns, but rather recognized several non-consensus sequences (Figure 5H). Next, we determined the relative enrichment of GGA consensus sites near the SRSF1-cross linked sites in the eCLIP dataset. Again, we observed an enrichment of GGA motifs near the SRSF1 crosslinking sites within the exons of control and MALAT1-depleted hypoxic cells (Figure 5I). However, we observed an overall reduction in the occurrence of GGA motifs near the SRSF1-cross linked sites within the introns of MALAT1-depleted hypoxic cells. These results imply that the observed increase in intronic binding of SRSF1 in MALAT1-depleted cells was due to sporadic non-consensus interaction of SRSF1 within the introns (Figure 5G-I). The change in SRSF1 binding was not due to the change in SRSF1 protein levels, as control and MALAT1-depleted cells showed comparable levels of total SRSF1 (Supplementary Figure 5D). Our RNA-seq showed reduced SRSF1 mRNA levels after 24 hrs of hypoxia. However, such changes in SRSF1 mRNA levels observed during 24 hrs of hypoxia did not correlate with SRSF1 protein levels, possibly due to inherent stability of SR-proteins. Overall, our results indicate that MALAT1 specifies the binding of SRSF1 to its strong consensus sites within target pre-mRNAs. Thus, in the absence of MALAT1, SRSF1 binds to the consensus sites (strong sites) in the exonic regions but in addition also recognizes the non-consensus sequences (weak sites). It is possible that ∼15000 copies of MALAT1 in hypoxic cells may potentially titrate several hundred thousand molecules of SRSF1. However, in the absence of MALAT1, these free pool of SRSF1 may start to recognize additional weak sites.

Our results revealed that most of the genes whose hypoxia responsive AS were modulated by the SRSF1/MALAT1 axis, were also localized near NSs. Both MALAT1 and SRSF1 are co-enriched in NSs^50,67^. We therefore examined whether the NS proximity of hypoxia-responsive genes influences the MALAT1-dependent recognition of SRSF1 to its substrate transcripts. To do this, we estimated the SRSF1 binding (average SRSF1 eCLIP peak number/transcript) to exons and introns of the top 1000 expressed speckle-near and speckle-far genes (gene relative position was determined based on SON-TSA-seq) in control and MALAT1-depleted hypoxic MCF7 cells. In control hypoxic cells, SRSF1 showed increased overall binding to transcripts of speckle proximal genes (Figure 5J). However, MALAT1 depletion resulted in overall decrease in the binding of SRSF1 to both speckle-near and -far genes (Figure 5J). This is in accordance with our earlier observation that MALAT1-depleted cells showed reduced binding of SRSF1 to exons (Figure 5G). In control hypoxic cells, intronic regions of speckle-near and -far genes showed reduced SRSF1 binding, as reflected by low eCLIP peaks/transcript (Figure 5K). However, in the MALAT1-depleted hypoxic cells, SRSF1 eCLIP peak/transcript were significantly increased in the introns of speckle-near, but not speckle-far genes (Figure 5K). These results imply that MALAT1 provides specificity to SRSF1-target mRNA interactions, preferentially to transcripts that are localized near NSs.

### MALAT1 enhances driving forces for SRSF1 condensate formation

In normoxic MCF7 cells, SRSF1-bound and MALAT1- and SRSF1-dependent hypoxia-responsive AS genes remain localized near NSs, and did not show significant movement towards NSs post-hypoxia (Figure 6A-6B). In addition, while MALAT1-depletion altered hypoxia-responsive AS, it did not change the positions of genes near NSs, again suggesting that MALAT1 does not influence the positioning of hypoxia-responsive genes near NSs (Figure 6A-B).

**Figure 6.**
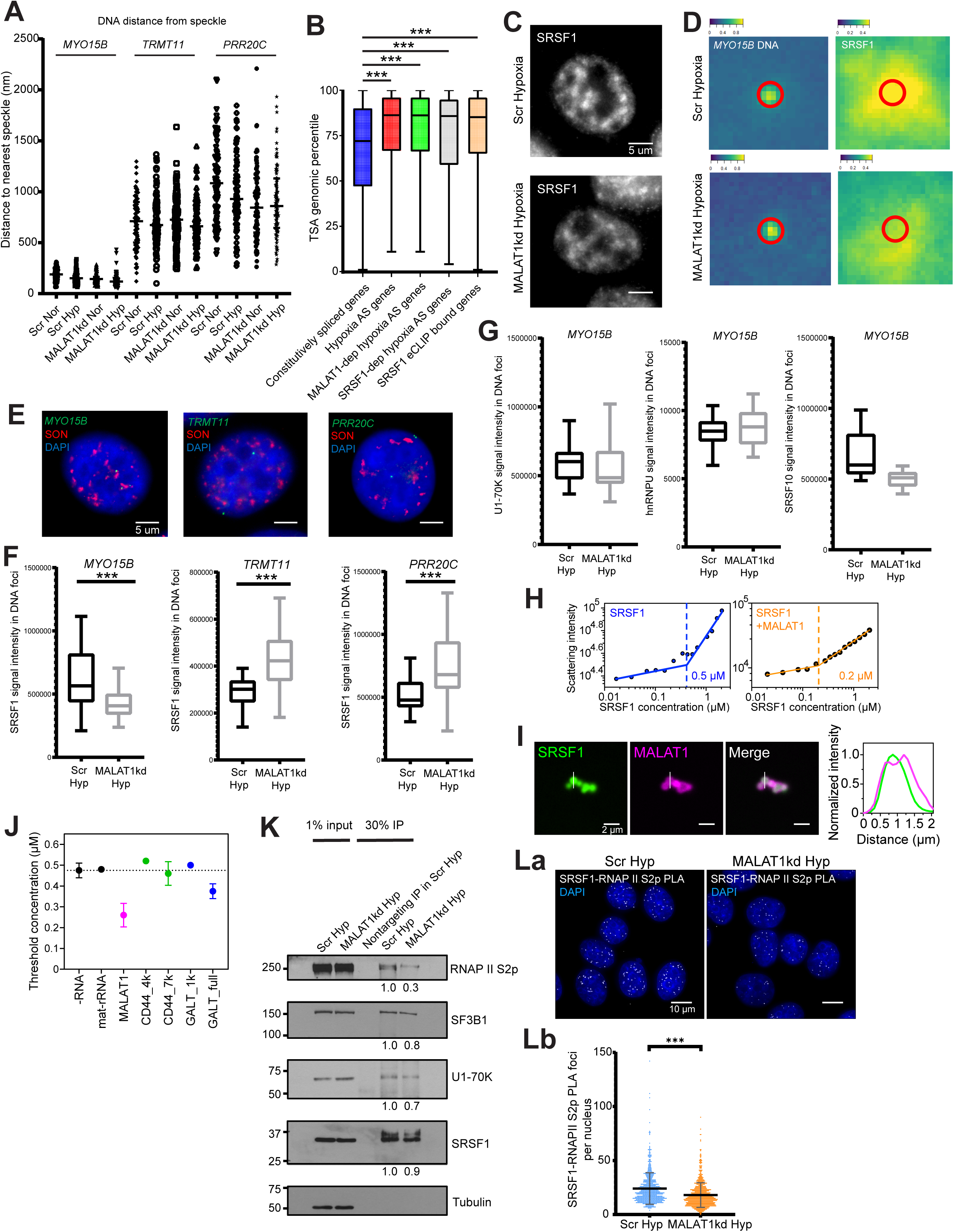
MALAT1 promotes SRSF1 condensate assembly. (A) Scatter plot showing quantification of distance between DNA FISH foci to the nearest NSs for genes in different SON-TSA percentiles. Error bars indicating mean and standard deviation. (B) Box plot showing TSA percentiles of all constitutively spliced genes, hypoxia-responsive AS genes, MALAT1- and SRSF-dependent AS genes, and SRSF1-bound RNA host genes. (C) SRSF1 immunostaining in control and MALAT1-depleted hypoxic MCF7cells. (D) Average intensity heatmap of SRSF1 immunostaining signal on or near DNA FISH signal. Grayscale values of the DNA FISH signal and SRSF1 immunostaining signal were converted to a matrix and the average intensity heatmap was calculated based on three biological replicates. (E) Representative image of DNA FISH and SON immunostaining for speckle-proximal MALAT1-regulated gene, speckle-far MALAT1-regulated gene, and non-hypoxia-responsive genes. (F) Quantification of SRSF1 signal intensity on DNA foci in control and MALAT1-depleted hypoxic MCF7. Box (Figures B, F, G) representing 25th to 75th percentile with line indicating mean value. Whiskers representing the min and max value. (*) P ≤ 0.05, (**) P ≤ 0.01, (***) P ≤ 0.001 by two-tailed student’s t-test. (G) Quantification of immunostaining signal of nuclear proteins intensity on speckle-proximal MALAT1-regulated gene DNA foci in control and MALAT1-depleted hypoxic cells. (H) 90-degree light scattering experiment of purified SRSF1 alone and with the presence of 1 nM MALAT1. (I) Confocal image of SRSF1-MALAT1 condensates. (J) Threshold concentrations of SRSF1 condensate formation without RNA (-RNA) and with the presence of 1 nM of mature rRNA, MALAT1, CD44_4kb, CD44_7kb, GALT 1kb, and GALT full length RNA as measured by 90 degree light scattering assay. (K) SRSF1 immunoprecipitation followed by immuoblot assays to detect various proteins in control and MALAT1-depleted hypoxic MCF7 samples. (L) Proximity ligation assay (PLA) of SRSF1 and RNAP II phosphor-Ser2 in control hypoxic cells and MALAT1kd hypoxic MCF7 cells. (*) P ≤ 0.05, (**) P ≤ 0.01, (***) P ≤ 0.001 by two-tailed student’s t-test. DNA is counter-stained with DAPI (blue; Fig E & La). See also Supplementary Figure 6.

Based on our results thus far, we hypothesize that MALAT1 influences the local concentration of SRSF1 at NS proximity, thereby providing specificity to the interactions between SRSF1 and substrate pre-mRNAs of NS-associated genes. More diffused nuclear localization of SRSF1 in MALAT1-depleted hypoxic cells supported this model (Figure 6C and Supplementary Figure 6A)^50^. Recent studies suggest that RNA polymerase II-mediated enrichment of splicing factors, including SRSFs on genes in the form of “splicing condensates” coordinates co-transcriptional pre-mRNA processing^68^. To determine whether MALAT1 influences local enrichment of SRSFs near hypoxia-responsive genes, we quantified the association of splicing factors, including SRSF1, on several hypoxia-responsive gene loci, which were prepositioned near or far from NSs in control and MALAT1-depleted hypoxic MCF7 cells (Figures 6D-G). The average intensity heatmap of SRSF1 IF (immunofluorescence) signal intensity from 50x50 pixel areas centered on the speckle-proximal DNA FISH gene foci showed that there was a decrease of SRSF1 IF signal on the gene locus upon MALAT1 depletion (Figure 6D). Additionally, the IF signal intensities of SRSF1 and other proteins were measured from 10-pixel diameter circular areas centered on the DNA FISH gene foci (Figure 6F-G). The hypoxia-responsive gene (*MYO15B*) located near NSs showed significantly decreased SRSF1 signal upon MALAT1 depletion (Figure 6E-F). Furthermore, MALAT1 along with SRSF1 was co-enriched near the MALAT1/SRSF1-dependent hypoxia-responsive AS genes at speckle proximity (Supplementary Figure 6B). However, hypoxia-responsive speckle-far (*TRMT11*) and non-hypoxia-responsive speckle-far (*PRR20C*) gene loci showed enhanced SRSF1 signal upon MALAT1 depletion, possibly due to the overall increase in SRSF1 diffuse nucleoplasmic pool in MALAT1 depleted cells (Figure 6E-F). Unlike SRSF1, other nuclear speckle-enriched splicing factors that we tested (U1-70K, hnRNP U and SRSF10) did not show significant changes in their association with the above-mentioned gene loci upon MALAT1 depletion (Figure 6G). Our results suggest that MALAT1-dependent enrichment of SRSF1 in NSs influences the enhanced association of SRSF1 to NS-proximal genes, contributing to hypoxia-responsive AS.

NSs are biomolecular condensates, which coordinate protein-protein and -RNA complex assembly for efficient pre-mRNA processing^69,70^. LncRNAs facilitate phase separation of proteins at various sub-cellular domains^71-75^. We hypothesize that MALAT1 modulates the local concentration of SRSF1 at NSs by influencing SRSF1 condensate formation. We first investigated the phase-separation of SRSF1 *in vitro* in the absence and presence of MALAT1 under identical solutions. We performed right-angle light scattering assays^76,77^ with purified SRSF1 over a 100-fold range in concentration where we observed a discontinuity in scattering intensity at ∼0.5 µM SRSF1, indicating the presence of a threshold concentration for the formation of higher-order assemblies (Figure 6H). We then used a mixture of fluorescently labeled SRSF1 and unlabeled SRSF1 and imaged the assemblies that form over a range of concentrations using confocal microscopy (Supplementary Figure 6C). Purified SRSF1 formed condensates at concentrations at or higher than 0.5 µM, indicating this to be the threshold concentration for phase separation of SRSF1 alone (Supplementary Figure 6C). In vitro droplet assays using fluorescently labeled SRSF1 and full-length MALAT1 RNA revealed that MALAT1 co-localized within SRSF1 condensates (Figure 6I). Most importantly, we observed that the presence of 5 x 10^-4^ molar ratio of MALAT1-to-SRSF1 reduced the threshold concentration for SRSF1 to form condensates by a factor of 2.5 (∼0.5 µM to ∼0.2 µM; Figure 6H & J and Supplementary Figure 6D). The change in the threshold concentration for phase separation can be explained using the formalism of polyphasic linkage^78,79^. Polyphasic linkage describes the influence of ligand (MALAT1) binding on the phase behavior of the macromolecule (SRSF1). MALAT1 can bind to SRSF1 in either the dense or dilute phase. If MALAT1 binding to SRSF1 is equivalent in the two phases, then the threshold concentration will remain unchanged (Equation 1 in STAR Methods). However, if MALAT1 preferentially binds to SRSF1 in the dense phase, then threshold concentration will shift down when compared to the intrinsic threshold concentration. Consequently, MALAT1 stabilizes the dense phase in the form of SRSF1 condensates. The converse will be true, and the threshold concentration will increase if the ligand, MALAT1, binds preferentially to SRSF1 in the dilute phase. Thus, based on the observed linkage between binding and phase equilibria, which shows that the threshold concentration is lowered by a factor of 2.5, we conclude that MALAT1 promotes the formation of SRSF1 condensates by binding preferentially to SRSF1 in the dense phase when compared to SRSF1 in the dilute phase.

To determine whether the effect on SRSF1 condensate formation is specific to MALAT1, we also measured SRSF1 threshold concentrations in the presence of several RNAs, including mature rRNA (mat-rRNA) and other SRSF1 substrate RNAs, including CD44 and GALT (Figure 6J and Supplementary Figure 6E-F). Two different lengths of CD44 and GALT were each produced by in vitro transcription, termed GALT (892 nts; 7 SRSF1 motifs), CD44 (4449 nts; 40 SRSF1 motifs), GALT_full (4361 nts; 57 SRSF1 motifs), and CD44_7k (7213 nts; SRSF1 78 motifs), encompassing different lengths and number of SRSF1 binding motifs within the RNA sequences. Light scattering assays revealed that none of these transcripts had affected the threshold concentration of SRSF1, though all the SRSF1 substrate transcripts localized within SRSF1 condensates (Figure 6J and supplementary Figure 6F). These results indicate that MALAT1 promotes SRSF1 condensate formation by binding preferentially to SRSF1 in the dense phase.

We observed that SRSF1 substrate mRNAs, such as CD44_7k co-localized with SRSF1 condensates (Supplementary Figure 6F). To evaluate if CD44_7k RNA competes with MALAT1 to be part of SRSF1 condensates, we performed measurements using a mixture of unlabeled SRSF1, a mixture of unlabeled and AlexaFluor 488-labeled MALAT1 (green), and a mixture of unlabeled and AlexaFluor 647-labeled CD44_7k mRNA (magenta) under different orders of addition (Supplementary Figure 6G). In the absence of SRSF1, RNA molecules alone failed to form condensates (Supplementary Figure 6G, top row, left). Upon addition of SRSF1, we observed SRSF1 condensates, containing higher MALAT1 concentrations compared to CD44_7k RNA (top row, right). In a pre-mixed sample of SRSF1+CD44_7k RNA, we observed condensate formation with fluorescence signals from CD44_7k RNA (middle row, left). Upon addition of MALAT1 (green) to the SRSF1+CD44_7k RNA mixture, MALAT1 displayed higher intensity compared to CD44_7k RNA (Supplementary Figure 6G, middle row right). In contrast, when CD44_7k RNA was added to the pre-mixed sample of SRSF1+MALAT1, CD44_7k RNA was unable to co-localize with the SRSF1 condensates, supporting the inference that MALAT1 shows higher affinity to SRSF1 condensates than CD44_7k RNA (Supplementary Figure 6G, bottom row). These results together suggest that MALAT1 enhances the driving forces for SRSF1 phase separation by binding preferentially to SRSF1 in the dense phase and thereby reducing the threshold concentration of SRSF1 required for phase separation. Furthermore, MALAT1 displays higher affinity to SRSF1 condensates over other pre-mRNAs, including several of the tested SRSF1 substrates.

### MALAT1 promotes the RNA pol II-SRSF1 splicing condensate assembly

We determined whether enhanced driving forces for SRSF1 phase separation by binding of MALAT1 influence inter-molecular interactions between SRSF1 and its binding partners. Condensates orchestrate molecular functions by influencing inter- as well as intra-molecular interactions^80-82^. Co-immunoprecipitation experiments revealed that MALAT1 enhanced the interaction between SRSF1 and several of its interactors (Figure 6K). For example, MALAT1-depleted cells showed significantly reduced interaction between SRSF1, and serine-2 phosphorylated elongation competent form of RNA polymerase II (Ser2p RNAP II), and small but consistent changes in the interaction with SF3B1 (U2 snRNP component) and U1-70K (U1 snRNP component) (Figure 6K). We further confirmed the involvement of MALAT1 in enhancing the intra-nuclear binding between SRSF1 and Ser2p RNAP II using a proximity ligation assay. MALAT1-depleted intact hypoxic MCF7 cell nuclei showed reduced SRSF1-Ser2p RNAP II interaction (Figure 6La-b). Ser2p RNAP II recruits SR-proteins to the actively transcribed genes^83-87^. Exon-bound SRSF1 stabilizes the U1-snRNP and U2 snRNP complexes at the splice sites as part of exon-definition^88-90^. In summary, MALAT1 facilitates the interaction between SRSF1 and Ser2p RNAP II, and snRNP components, and defects in such molecular interactions would affect the recruitment of SRSF1 and core spliceosome components to the RNA pol II-engaged actively transcribing genes.

Recent studies revealed that Ser2p RNAP II dictates the formation of SRSF-containing “splicing condensates” on genes for efficient co-transcriptional splicing^68^. We hypothesize that MALAT1 modulates AS by influencing RNAP II-mediated SRSF splicing condensate assembly. We tested whether MALAT1 facilitated intermolecular interactions between SRSF1 and the CTD of RNA pol II by promoting the formation of SRSF1 condensates. The carboxy-terminal part of the CTD repeat (CTD repeats 27-52) supported co-transcriptional processing, including splicing, whereas the amino-terminus CTD (CTD repeats 1-20) supported only 5’-pre-mRNA capping^91^. We therefore used the CTD repeats 27-52 of RNA pol II, instead of full-length CTD (CTD repeats 1-52) at concentrations, which do not undergo phase separation on its own (Supplementary Figure 7A), to test the potential involvement of CTD in promoting SRSF1 condensate formation. We used both unphosphorylated and ser-2 phosphorylated (in vitro phosphorylated by CDK9/Cyclin T) forms of CTD (CTD and pCTD). By confocal microscopy, we observed that both CTD and pCTD co-localized with SRSF1 condensates (Figure 7A and Supplementary Figure 7B). We next determined whether CTD and pCTD influence SRSF1 assembly by measuring threshold concentrations for phase separation by right angle light scattering for a mixture of equimolar CTD or pCTD and SRSF1. We observed that the SRSF1 threshold concentration was lowered from ∼0.5 µM to ∼0.2 µM in the presence of unphosphorylated pol II CTD, to the same extent of the effect observed in the presence of MALAT1 (5 x 10^-4^ molar ratio of MALAT1-to-SRSF1) (Figure 6H and 7B). Strikingly, Phosphorylated-CTD (pCTD) further reduced the SRSF1 threshold concentration to form condensates to ∼0.1 µM (Figure 7B). These results indicate that CTDs, like MALAT1, enhances the driving forces for the formation of SRSF1 condensates and preferentially binds to the SRSF1 condensates over the free pool of SRSF1. Furthermore, the pCTD even more efficiently promotes SRSF1 condensation (∼5-fold compared to SRSF1 alone, and ∼2-fold compared to CTD- or MALAT1-containing SRSF1).

**Figure 7.**
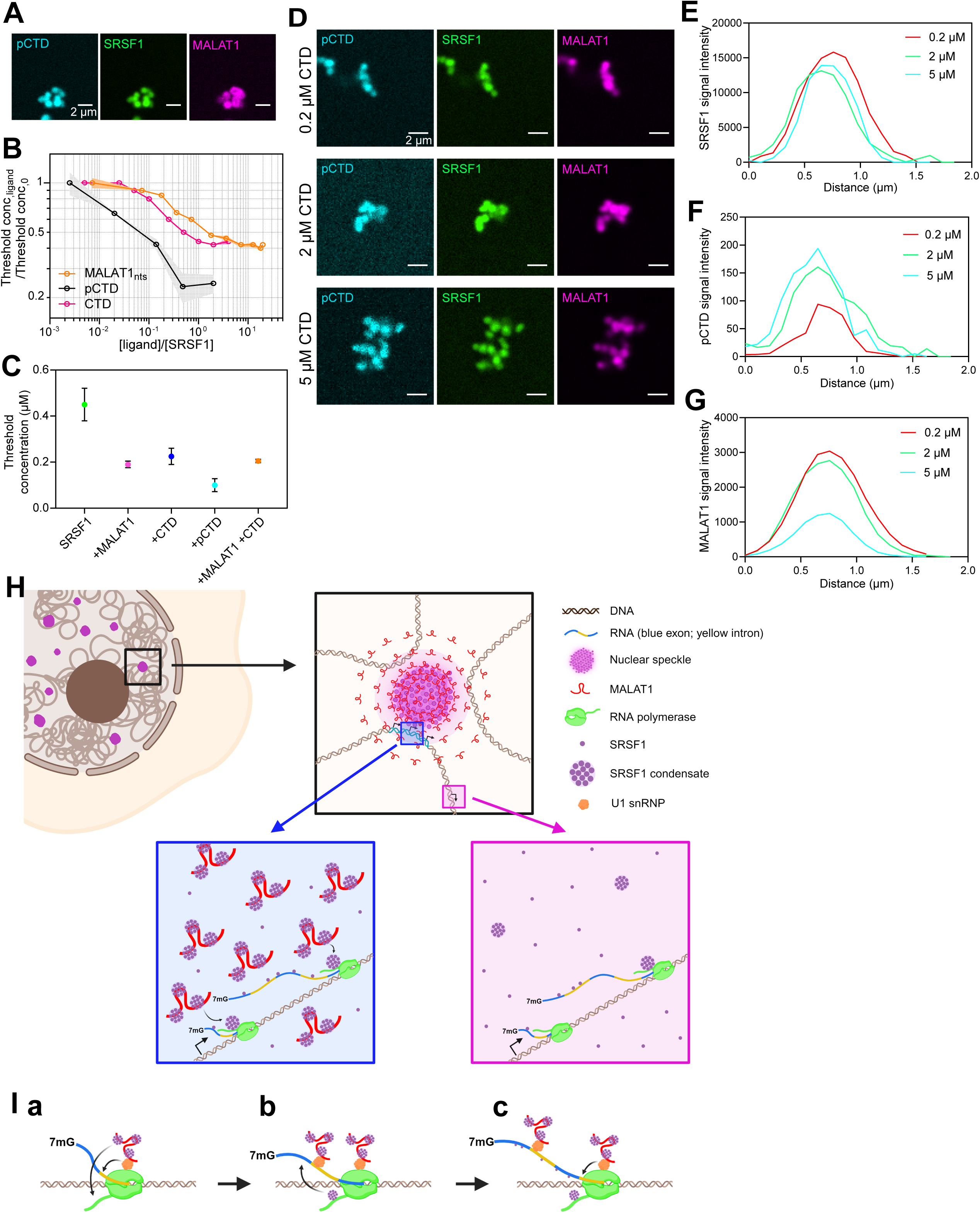
MALAT1 promotes the RNA pol II-SRSF1 splicing condensate assembly. (A) Confocal image of SRSF1-pCTD-MALAT1 condensates. (B) Graph showing the change in the SRSF1 threshold concentration over change in concentration of MALAT1, CTD, and pCTD. (C) Threshold concentrations of SRSF1 condensate formation in presence of MALAT1, CTD, pCTD, or MALAT1 + CTD as measured by 90 degree light scattering assay. (D) Confocal images of SRSF1 condensates (green) incubated with MALAT1 (pink) and increasing concentrations of pCTD (cyan). (E-G) Line profile showing SRSF1 (E), pCTD (F), and MALAT1 (G) signal intensities in the condensates with increasing concentrations of pCTD. (H) Model showing hypoxia-responsive genes pre-positioned near to a nuclear speckle. In the speckle-proximal region, MALAT1 RNA facilitates the formation of SRSF1 condensates and promotes the association of SRSF1 to RNAP II and the pre-mRNA. (I) Schematic showing how MALAT1 RNA, recruited to transcriptionally active genes via U1 snRNP, promotes the formation of SRSF1 condensates, which then interact with elongating RNA pol II and the pre-mRNA. See also Supplementary Figure 7.

To compare the effect of MALAT1 and CTDs (CTD and pCTD) on SRSF1 condensate formation, we measured threshold concentrations at multiple molar ratios of SRSF1:MALAT1 and SRSF1: CTD or pCTD. Interestingly, we observed that CTD and pCTD further reduced the threshold concentration for SRSF1 phase separation compared to the same molar ratio of MALAT1 to SRSF1 (Figure 7C). We observed consistent results by microscopy where we co-localized of SRSF1, MALAT1, and CTD (or pCTD) in mixtures with different orders of addition. First, we generated SRSF1 condensates by incubating 1 µM SRSF1 with 1 nM MALAT1. Then, we added CTD or pCTD over a range of concentrations and determined the relative concentrations of all three molecules within the condensates by quantifying their fluorescence intensities (Figure 7D-G and Supplementary Figure 7B-E). SRSF1 did not show significant change in its overall concentrations (Figure 7D-E and Supplementary Figure 7B-C). On the other hand, we observed an inverse correlation between MALAT1 and CTD or pCTD intensities within the SRSF1 condensates (Figure 7D-G and Supplementary Figure 7B-E). SRSF1 incubated with higher concentrations of CTD or pCTD contained lower levels of MALAT1 (Figure 7D & G and Supplementary Figure 7B & E). Next, we changed the order of addition and added MALAT into the SRSF1-CTD pre-formed condensates. Indeed, MALAT1 did not efficiently load into the pre-formed CTD-containing SRSF1 condensates, suggesting that CTD and pCTD displaces MALAT1 from the SRSF1 condensates (Supplementary Figure 7F). Based on these results, we propose that MALAT1 initially promotes the formation of SRSF1 condensates by reducing the threshold concentration required for forming the phase condensates. RNA pol II preferentially interacts with SRSF1 condensates over the free pool of SRSF1. MALAT1-promoted SRSF1 condensates are recognized and are efficiently transferred to RNA pol II CTD for generating splicing condensates at the transcription sites.

## Discussion

Hypoxia-responsive gene expression is studied extensively, especially due to its relevance in tumor biology, such as angiogenesis, EMT, and chemoresistance^1,2^. Besides transcription, cancer cells, in response to hypoxia, achieve differential gene expression by modulating AS of ∼1000 pre-mRNAs without altering their overall transcription^6,12,21,92^. Hypoxia-specific isoforms of proteins control processes such as angiogenesis and cancer metastasis^10^. However, very little is known about the underlying mechanism/s controlling hypoxia-responsive AS. In this study, we unraveled the role of higher-order genome, nuclear domain organization, and a chromatin-associated lncRNA, MALAT1, in hypoxia-responsive AS.

Transcriptionally active genes preferentially localize at NS proximity^22,32,93^. Also, genes localized in the NS neighborhood showed enhanced transcription and RNA processing^15,22,28,33-35^. For example, heat shock-responsive genes were either pre-positioned or moved close to NSs upon heat shock to amplify their expression^34^. During ribotoxic stress, re-localization of several immediate early genes near to NSs facilitated their transcription and co-transcriptional splicing^35^. Finally, in response to DNA damage, ∼20% of p53 target genes, repositioned to NSs for hyperactivation^33^. These results imply the role of NSs in boosting stress-induced gene activation. Contrary to these observations, most genes induced in response to 24 hrs of acute hypoxia did not move towards NSs. Instead, they were pre-positioned close to NSs, even in normoxic cells. Hypoxia, in turn, enhanced their expression or AS, without significantly altering the gene position. Even the ∼100 HIF (HIF1 and 2) target genes, which displayed hypoxia-responsive induction in MCF7 cells, were pre-positioned near NSs (mostly within the SPADs) in normoxic cells. On the other hand, the HIF target genes, which were not induced after hypoxia in MCF7 cells, tend to position in speckle distal regions. These results imply that NSs boost the expression of genes that are in its proximity. We analyzed the status of gene position post-24hrs of hypoxia. In the p53 study, authors observed NS association of some of the genes only during specific time windows post DNA damage^33^. We might be able to detect NS proximity repositioning of a few or more genes if we analyze the data after several time points of hypoxia.

A recent study reported the potential involvement of HIF2-α in facilitating the repositioning of ∼25% of the HIF2-α target genes to speckle proximity for their activation^94^. In clear cell renal carcinoma (ccRCC) cells exhibiting constitutive activation of HIF2-α, the authors demonstrated that HIF2-α promoted the physical association of a subset of HIF2-target genes to speckles^94^. Surprisingly, in MCF7 cells, we did not observe such positive correlation between hypoxia-induced repositioning of HIF2-α target genes to NSs and trascription activation. The discrepancy in results between these two studies could be due to different cell types (BrCa versus ccRCC cells) and/or the method used to induce hypoxia. We exposed MCF7 to acute hypoxia (0.2% oxygen for 24 hours) and the study utilizing ccRCC cells used cells with an inactivating mutation within pVHL, which is an inhibitor of HIFs, thus making the HIFs constitutively active^95^.

We observed that genes undergoing hypoxia-responsive AS are also pre-positioned near NSs compared to hypoxia non-responsive constitutively spliced genes. Since, NSs are enriched with RNA binding proteins (RBPs), including splicing factors, NS-associated genes show increased association of spliceosomes, resulting in enhanced splicing efficiency^28,96^. Even in-silico modeling studies revealed that a slight increase in splicing protein localization into NSs leads to a profound enhancement in splicing activity^97^. These observations indicate that the distance between genes and NSs and the enrichment of spliceosome components near genes dictate splicing efficiency. We hypothesize that the pre-positioning of hypoxia-responsive genes near NSs in normoxic cells promote the hypoxia-induced AS due to enhanced interaction between their transcripts and speckle-resident splicing factors.

We propose that MALAT1 promotes hypoxia-responsive AS of genes located near NS by regulating splicing factor activity. Several pieces of data support our model. MALAT1 is enriched at the periphery or the “shell” of NS, where actively transcribed genes, including hypoxia-induced genes, are distributed^31^. MALAT1 is shown to interact with active and differentially spliced RNA pol II-transcribed genes^54,55,98-100^. Such chromatin and NS association of MALAT1 is dependent on active transcription and its interaction with NS-enriched spliceosome components such as U1-snRNPs^55,101^. In addition, MALAT1 binds to dozens of NS-resident RNA splicing factors, such as SRSFs^50,56-58^. For example, hypoxic cells harbor ∼15,000 copies of MALAT1/nucleus, and each MALAT1 could potentially interact with ∼100s SRSF1. This suggests that MALAT1 by enhancing the local concentration of SRSF1 near hypoxia-responsive genes that are pre-positioned proximal to NSs, influences the molecular interactions between SRSF1 and target pre-mRNAs. In support of this model, we demonstrated that SRSF1 co-regulated the AS of ∼31% of MALAT1-dependent hypoxia-responsive AS. Further, SRSF1 was identified as the top target protein, binding to MALAT1-regulated alternatively spliced exon junctions within NS-resident genes. Finally, the altered binding of SRSF1 to NS-associated pre-mRNA and its reduced association to NSs and NS resident hypoxia-responsive gene loci upon MALAT1 depletion further confirm the role of MALAT1 in modulating SRSF1 function.

The question that we tried to address here is how MALAT1 regulates SRSF1 activity. The traditional model of co-transcriptional splicing describes the role for the elongation-competent ser2 CTD phosphorylated form of RNAP II (Ser2p RNAP II), in recruiting SRSFs to pre-mRNAs^83,85,86,102-105^. However, this direct recruitment model alone fails to explain the AS of only a sub-set exons in response to various cellular stimuli. Firstly, a single CTD within the RNA pol II cannot activate simultaneous recruitment of various RNA processing factors to pre-mRNAs^70^. Also, transient interactions between elongating RNAP II and SRSFs at the gene locus merely by random interaction will be inefficient. Instead, high-efficiency molecular interactions could be achieved by localized enrichment of SRSFs near transcription sites. We argue that MALAT1 promotes RNA pol II-mediated recruitment of SRSFs to pre-mRNAs by maintaining not only the critical concentration of SRSFs at the transcription sites of NS-associated genes, but also in a form as SRSF1 condensates, which is preferentially recognized RNA pol II, (Figure 7H). Condensates enhance the local concentrations of molecules in defined areas, such as NDs, and orchestrate intermolecular interactions^16,80,81^. Based on our results, we propose a model in which MALAT1 enhances the driving forces for SRSF1 condensation at NS proximity by reducing the threshold concentration (from 0.5 to 0.2 µM) required to form the SRSF1 condensates. By this, MALAT1 increases the local concentration of SRSF1 condensates near NSs (Figure 7H-I). RNA pol II CTD, especially Ser2p CTD, preferentially interacts with the MALAT1-initiated SRSF1 condensates over diluted phase for efficient co-transcriptional splicing. Based on our results, we argue that MALAT1 promotes the Ser2p RNA pol II-mediated SRSF containing “splicing condensates” on active transcription sites for efficient co-transcriptional splicing^68^. Finally, we demonstrate that RNA pol II pCTD displaces MALAT1 from SRSF1 condensates. These observations partially explain a mechanism in which at the transcription sites, the phase equilibria shift toward condensates being enriched in Ser2p RNA pol II, implying a displacement of MALAT1 from SRSF1 condensates.

Several earlier studies have demonstrated that elongating competent RNA pol II interacts and recruits U1 snRNP to actively transcribing genes^106,107^. MALAT1 is recruited to the transcriptionally active chromatin interacting with U1 snRNP^55,101^. Based these, we propose that MALAT1 is initially recruited to the actively transcribing genes, especially near splice junctions, by RNA pol II-bound U1 snRNP (Figure 7I). MALAT1 by binding to a myriad of RBP, such as SRSF1, preferably by promoting the formation of RBP condensates, increases the local concentration of these factors near the transcribing gene (Figure 7I). Transcription-engaged RNA pol II preferentially recognizes the MALAT1-associated splicing factor condensates and recruits them to pre-mRNA to facilitate co-transcriptional splicing (Figure 7I). In summary, we provided insights into the molecular function of MALAT1 that it promotes the formation of SRSF1 condensates near NSs. This allows the SRSF1 to reach a local threshold concentration that facilitates intermolecular interactions, such as the formation of Ser2p RNA pol II-mediated SRSF1-containing splicing condensates at transcription sites.

## Acknowledgements

We thank members of the Prasanth lab for their valuable comments. We thank Dr. Liguo Zhang and Pradeep Kumar (Belmont lab) for their guidance in the SON TSA-seq experiment and data analyses. We thank Dr. Kevin Van Bortle (UIUC) for the scientific discussions. We thank Dr. Benita Katzenellenbogen (UIUC) for MCF7 cells; Dr. Matthew King and Avnika Pant (WashU) for cloning SRSF1 and preparing purified proteins and RNA substrates. We thank Dr. Brian Freeman (UIUC) and Dr. Janhavi Kohle for technical help. This work was supported by grants from the National Institutes of Health T32EB019944 to YJS and SB, R21-AG065748 & R01-GM132458 to KVP, GM125196 to SGP, K99GM152778 to MKS, R01-HL126845 to AK, Muscular Dystrophy Association Research Grant MDA1072487 and Chan-Zuckerberg Biohub Chicago Awards to AK, the St. Jude Collaborative on the Biophysics and Biology of RNP granules to RVP, Cancer center at Illinois seed grants and Prairie Dragon Paddlers to KVP and AK, ARPA-H (KVP), and National Science Foundation (NSF) (KVP [1723008 EAGER] & 2243257 NSF center for quantitative cell biology).

## Methods

### Cell lines

MCF7 and MDA-MB-231 cells were grown in RPMI 1640 medium. HEK293T cells were grown in DMEM medium. MDA-MB-231 and HEK293T media were supplemented with 10% fetal bovine serum and penicillin/streptomycin. MCF7 medium was supplemented with 5% fetal bovine serum and penicillin/streptomycin. Cells were maintained in a 5% CO2 incubator at 37 degrees. For hypoxia treatment, cells were incubated in a 0.2% O2 and 5% CO2 hypoxia chamber for 24 hours. We confirm that the identity of all cell lines used in our study has been authenticated by STR profiling. All cell lines were regularly checked for mycoplasma using the ATCC Universal Mycoplasma Detection Kit.

### Generation of MALAT1 CRISPR KO cell lines

MALAT1 CRISPR KO clones were made by transiently transfecting spCas9, MALAT1 gRNA, AB59, CMV-Puro, and CMV-Blast in MCF7, MDA-MB-231, and HEK293T cells. Selection was carried out with 2 μg/ml of puromycin and 10 ug/mL blasticidin followed by single clone selection. The KO clones were confirmed by real time PCR.

### Knockdown by ASO and shRNA treatment

Knockdown of MALAT1 was done in two rounds of ASO treatment at a final concentration of 200 nM and 100 nM using Lipofectamine RNAiMax reagent (Invitrogen). After the second round of knockdown, the cells were cultured for 24 hours in normoxia or hypoxia before collection. Knockdown of SRSF1 was carried out by preparing shRNA-expressing lentivirus in HEK293T cells transfected pPAX2, pMD2.G, and a pLK0.1 plasmid containing an shRNA against SRSF1 and Lipofectamine3000.

### RNA extraction and quantitative real-time PCR (RT-qPCR)

RNA was extracted using Trizol reagent (Invitrogen) as per manufacturer’s instructions. Samples for RNA-seq were further cleaned up by RNeasy Mini Kit (QIAGEN). RNA was reverse transcribed into cDNA by Multiscribe Reverse Transcriptase and Random Hexamers (Applied Biosystems). RT-qPCRs were performed using the StepOne Plus system (Applied Biosystems). Transcript levels were quantitated against a standard curve by Real-time RT-PCR using the SYBR Green I fluorogenic dye and data analyzed using the StepOne Plus system software. 18S rRNA is used as a reference gene to normalize gene expression.

### Bioinformatics and statistical analyses of RNA-seq data

The RNA-seq libraries were prepared with Illumina TruSeq Stranded mRNAseq Sample Prep kit (Illumina). Paired-end, polyA+ RNA-sequencing was performed on the Illumina Novaseq 6000 platform at the Roy J. Carver Biotechnology Center at UIUC. High quality of RNA-seq reads was confirmed by FastQC^1^. RNA-seq reads were aligned to the human reference genome GRCh38 assembly using STAR^2^. Gene annotation was based on the GRCh38 assembly GTF file downloaded from Ensemble (v98). For statistical analyses, raw gene counts were first analyzed by HTSeq-count^3^, then analyzed using edgeR^4^. Libraries were normalized using the TMM method. Differential expression analyses were performed using exactTest. Genes were considered as having significantly different expression following imposed cutoff clearance (FDR (q-value) <0.05, log2(fold change) >1). Differential splicing analysis was performed using rMATS (version 3.2.5), and significant events were identified with imposed cutoffs (FDR <0.05, junction read counts ≥10, PSI ≥15%)^5^. Heatmaps were created using the pheatmap package in R^6^, with row centering and scaling. The chromosome-wide heatmaps were drawn using the chromoMap package in R^7^. The color scale was based on the Z-score, with the red-blue color scale for differential expression and yellow-violet color scale used for differential splicing. The hypoxia-responsive DEGs and AS events were identified based on the comparison between the control normoxic and control hypoxic RNA-seq data, using the analysis and cutoffs mentioned above. The MALAT1-dependent hypoxia-responsive DEGs and AS events were identified by first determining the MALAT1-independent genes based on the comparison between MALAT1-depleted normoxic and MALAT1-depleted hypoxic RNA-seq data, wherein the DEGs and AS events in the MALAT1-depleted normoxic vs MALAT1-depleted hypoxic RNA-seq were determined to be MALAT1-independent hypoxia-responsive events. The hypoxia-responsive DEGs and AS events that were identified in the control samples but not in the MALAT1-depleted samples were determined to be MALAT1-dependent hypoxia-responsive events.

Gene ontology analyses (biological processes, Kegg pathway analyses) and GSEA (gene set enrichment analysis) were performed using clusterprofiler of Bioconductor^8^. Specifically, gene ontology for biological processes was performed using the enrichGO function, Kegg pathway analyses was performed using enrichKEGG. All enrichment analyses include using a background gene list containing all 24087 genes which showed qualifiable expression in the RNA-seq. GSEA analysis was performed using gseGO function and gene lists were ranked using logFC values. The enrichment of RBPs on the AS splicing regions (alternatively spliced exon plus 500 bp of the flanking introns) were calculated using MEME suite based on the cisBP-RNA database. The constitutively spliced regions were selected based on the splicing events identified by rMATS that were not identified as hypoxia-responsive, then randomly selecting an equal number of the constitutively spliced events to the MALAT1-dependent hypoxia-responsive cassette exon AS events to compare the enrichment of the RBPs.

### Transwell migration and invasion assay

Cells were grown in regular media until 50% confluency then the medium was changed to low serum medium (1% FBS in RPMI 1640) for 12 hours. Cells were harvested using trypsin and resuspended in low serum medium to a concentration of 2000 cells per uL for the migration assay and 5000 cells per uL for the invasion assay. 100 uL of the cell suspension was added to the upper chamber of the transwell and the lower chamber of the transwell was filled with 700 uL of the regular growth media. The transwell plate was then put in the incubator or the hypoxia chamber for 24 hours. The non invaded cells in the upper chamber of the transwell were scraped off thoroughly before fixing the cells and staining with 0.05% crystal violet in 10% methanol.

For the invasion assay, the matrigel-coated transwells were thawed to room temperature then incubated with low serum media at 37 degrees Celsius to rehydrate before seeding the cells.

### Scratch wound healing assay

Cells were grown in regular media until confluence, then the medium was changed to low serum medium (1% FBS in RPMI 1640) for 12 hours. Scratches were drawn using a sterile 10 uL pipette tip and the cells were rinsed twice with low serum medium to remove the debris and floating cells. Cells were allowed to migrate at 37 degrees Celsius in normoxic or hypoxic conditions and the wound area was imaged every 15 minutes. Total migrated cells was calculated using the 0 and 20 hour time points for each position. 5 positions were chosen per replicate and the experiment was performed in triplicates.

### Lactic acid secretion assay

MCF7 cells were seeded to 35 mm plates. MALAT1 knockdown and control scramble knockdown were performed. After quenching the knockdown by addition of media, the cells were grown for 48 hours and the lactic acid levels in the media were measured using the Promega Lactate-Glo Assay (Cat # J5021). The lactic acid levels in control and MALAT1 kd samples were measured from three biological replicates, and the values for each replicate was based on the average of three replicate wells.

### Mouse xenograft

All protocols involving animals were approved by the Institutional Animal Care and Use Committee (IACUC) at the University of Illinois Urbana-Champaign. Methods regarding tumor models have been previously described^9^. Briefly, MDA-MB-231 and HEK293T cells were cultured until 70% confluency and then grafted. For orthotopic tumors, cells were grafted into the mammary fat pad and followed through time by direct caliper measurement. For experimental metastasis, cells were grafted intravenously via the tail vein, and followed through time by bioluminescence imaging (IVIS).

### eCLIP-seq

SRSF1 eCLIP was performed in concordance with previously published protocols^10^. UV crosslinking was performed with 1 pulse of 400 mJ/cm2. Briefly, crosslinked cells were lysed in buffer and sonicated, followed by treatment with RNase I (Thermo Fisher Scientific) to fragment RNA. SRSF1 antibody (Bethyl Labs A302-052A) was pre-coupled to sheep anti-rabbit IgG Dynabeads (Thermo Fisher 11203D), added to lysate, and incubated 4h at 4°C. Prior to IP washes, 2% of sample was removed to serve as the paired input sample. For IP samples, high- and low-salt washes were performed, after which RNA was dephosphorylated with FastAP (Thermo Fisher Scientific) and T4 PNK (NEB) at low pH, and a 3′ RNA adaptor was ligated with T4 RNA ligase (NEB). 8% of IP and input samples were run on an analytical 4–12% PAGE gel, transferred to polyvinylidene fluoride membrane, blocked in 5% dry milk in tris-buffered saline with Tween, incubated with anti-SRSF1 antibody, washed, incubated with HRP-conjugated anti-rabbit secondary, and visualized with chemiluminescence imaging to validate successful IP. The remaining IP and input samples were run on a 4–12% PAGE gel and transferred to nitrocellulose membranes, after which the region from the protein size to 75 kDa (∼200 nt) above protein size was excised from the membrane, treated with proteinase K (NEB) to release RNA, and concentrated by column purification (Zymo).

Input samples were then dephosphorylated with FastAP (Thermo Fisher Scientific) and T4 PNK (NEB) at low pH, and a 3′ RNA adaptor was ligated with T4 RNA ligase (NEB) to synchronize with IP samples. Reverse transcription was then performed with AffinityScript (Agilent), followed by ExoSAP-IT (Affymetrix) treatment to remove unincorporated primer. RNA was then degraded by alkaline hydrolysis, and a 3′ DNA adaptor was ligated with T4 RNA ligase (NEB). qPCR was then used to determine the required amplification, followed by PCR with Q5 (NEB) and gel electrophoresis to size-select the final library. Libraries were sequenced on the NovaSeq6000 platform (Illumina). eCLIP was performed on IP from two independent samples along with paired size-matched input before the IP washes.

### eCLIP computational analysis

To analyze the eCLIP reads, the CLIP tool kit (CTK) pipeline was used^11^. Briefly, 3’ adaptors were clipped from reads, followed by collapsing of PCR duplicates and removal of the N10 random barcode sequence. Reads were then mapped using the bwa aligner and further cleaned up for PCR duplicates and alignments to repetitive and ribosomal non-coding RNA regions. Peak calling was performed on the remaining mapped reads as well crosslinking site analysis using CIMS/CITS. SRSF1 peaks were annotated to the nearest gene as well as the RNA feature, including intergenic, intron, exon-intron, and exon, wherein exon was further categorized into CDS and UTRs. Top 6-mers enriched in CLIP experiments were identified as detailed before in Feng et al.^12^ and analysis was changed to 6-mer instead of 7-mer.

### TSA-DNA-seq

MCF7 cells were seeded to T300 flasks and cultured at 37 degrees at 5% carbon dioxide for 24 hours. After the second round of ASO treatment, the cells were cultured in normoxic or hypoxic (0.2% oxygen) conditions for 24 hours. SON TSA-seq was carried out using the protocol for SON TSA-seq 2.0 on attached cells, with Condition E as described^13^ with a minor modification: cells were fixed by adding freshly made 8% PFA to the plates at a final concentration of 1.6% PFA. Libraries were constructed using Illumina TruSeq ChIP Sample Prep Kit and were amplified by 9-11 cycles of PCR. Libraries were sequenced on the NovaSeq6000 platform (Illumina).

### TSA-DNA-seq analysis

The TSA-DNA sequencing data were processed as previously described^13^. Briefly, we mapped raw sequencing reads to the human reference genome (hg38, Chromosome Y excluded) using Bowtie2 (version 2.1.0) with default parameters^14^. We applied the rmdup command from SAMtools (version 1.5) to remove potential PCR duplicates in the alignment^15^. We then used these alignment files as the input files for TSA-seq normalization. The smoothed TSA-seq enrichment scores were ranked over all 20-kb bins genome-wide from largest to smallest excluding hg38 unmapped regions, dividing these ranked scores into 100 equal sized groups, defined as percentiles 1 to 100 (lowest to highest scores). Adjacent 20-kb bins ranked in the 95th−100th percentile were merged to segment these regions as SPADs. To determine changes in TSA scores during hypoxia or MALAT1kd, the TSA enrichment scores were rescaled as previously described^13^ and a pairwise comparison was performed based on the rescaled TSA scores of the biological replicates where we calculated the P-value for the genomic bins then filtered for the regions that showed significant changes between the samples.

### DNA FISH and immunostaining

DNA FISH probes were made using Nick Translation Kit (Abbott Molecular) as per manufacturer’s instructions. Digoxin-labeled RNA probes were in vitro transcribed as per manufacturers’ instructions (DIG RNA labeling Mix, Roche; T7 polymerase, Promega; SP6 Polymerase, Promega) and purified by G-50 column (GE Healthcare). Cells on coverslips were fixed with 3% PFA then quenched with 0.5 M glycine. The cells were permeabilized with 0.5% Triton X-100 and blocked with 5% normal goat serum. Primary and secondary antibodies were diluted in 1% normal goat serum. Antibody incubations were done at room temperature for 1 hour. After secondary antibody incubation and washing, the coverslips were post-fixed with 3% PFA and quenched with 0.5 M glycine. The coverslips were treated with 0.1 mg/mL RNase A at 37 degrees for 1 hour and incubated with 0.1 N HCl for 10 minutes. The coverslips were washed with 2X SSC then incubated in 50% formamide in 2X SSC for 1 hour. DNA FISH probes were mixed with salmon sperm DNA, yeast tRNA, and human Cot-1 DNA and water, 20X SSC, formamide, and dextran sulfate were added to a final concentration of 2X SSC, 50% formamide, 10% dextran sulfate. The probe mix was added to the coverslip and heat denatured at 80 degrees for 5 minutes. The probes were then hybridized in a humidified chamber at 37 degrees overnight. The coverslips were then washed in 2X SSC, 0.1X SSC, and 4X SSC then mounted using a VectaShield mounting medium. For imaging, z-stack images were taken using DeltaVision microscope (GE Healthcare) equipped with 60 X/1.42 NA oil immersion objective (Olympus) and CoolSNAP-HQ2 camera. Images were processed using FIJI/ImageJ. For the average intensity heatmap, the positions of the regions of interest (ROIs) were selected based on the focus and position of the DNA FISH foci, wherein the z-position was determined on the focus of the DNA FISH foci and the 50x50 pixel area was centered on the foci. After choosing the ROIs, the signal from the immunostaining channel (e.g. for SRSF1, using the channel corresponding to the dye-conjugated secondary antibody) was extracted for each ROI. The ROI images were stacked and the average intensity was calculated using the image stack and converted to a matrix. The matrix values were then used to generate a heatmap, and the scale was kept the same for control and MALAT1-depleted samples. To calculate the amount of immunostaining signal on the DNA foci, the ROIs were selected based on the position of the DNA FISH foci in the DNA FISH IF images. The ROIs were based on a 10-pixel diameter circular area centered on the DNA FISH foci with the z-position based on the focus of the foci. The integrated signal intensity of the immunostaining channel was calculated for each ROI in control and MALAT1-depleted samples, then plotted as a box plot showing the mean, 25^th^ percentile, and 75^th^ percentile values.

### Immunoprecipitation

Cells were washed with PBS and lysed in lysis buffer (50 mM Tris pH 7.4, 100 mM NaCl, 5 mM MgCl2, 0.5% NP-40, 0.3% Triton X-100, 1 mM DTT, 0.5 mM EDTA, and 10% glycerol) containing phosphatase and protease inhibitors. Lysates were then sonicated and centrifuged at 10000 g for 10 min to remove insoluble debris. Next, lysates were pre-cleared with Gammabind G sepharose (GE Healthcare Life Science) for 30 min at 4 °C. Antibodies were then added into lysates and incubated at 4 °C overnight. Proteins bound by antibodies were pulled down by Gammabind G Sepharose for two hours at 4°C. After incubation, beads were washed in lysis buffer and captured proteins were eluted and analyzed with western blot.

### Generation of DNA constructs

pGEMT-MALAT1 was cloned by amplifying the MALAT1 cDNA from pCMV-MALAT1 and ligating it into the pGEM-T easy vector (Promega A1360) according to the manufacturer’s protocol. pET28-SRSF1 was cloned by subcloning SRSF1 cDNA from YFP-SRSF1 into the pET28 vector by restriction digestion and ligation. The pET28-SRSF1 was transformed into the BL21-CodonPlus(DE3)-RIPL cells as described below.

Total genomic DNA was isolated using the Wizard® Genomic DNA Purification Kit (Promega, USA) as follows. Approximately 5 million HEK293T cells were harvested and washed once with PBS. Cells were lysed in 600µL lysis buffer by vortexing followed by a 5 minute incubation with protein precipitation buffer. Lysates were centrifuged at 14,000 rpm for 4 minutes, and the supernatant was transferred to a tube with 600µL isopropanol. After a 1 minute centrifugation at 14,000 rpm, the pellet was washed with 70% ethanol then left to dry by air for ∼15 minutes. After resuspension in 100µL ddH2O, the DNA concentration and purity were measured by Nanodrop confirmed the ratio of absorbances at 260nm and 280nm to be ∼2.0, indicating pure DNA. This genomic DNA was used as a template to PCR amplify the GALT_1k, GALT_full, or CD44_4k, and CD44_7k genes using custom primers. Each PCR product was confirmed to be a single DNA product with the correct molecular wight by gel electrophoresis. Then, 1 µL of PCR product was directly mixed with 1µL of pCR™Blunt II-TOPO™ provided in the Zero Blunt™ TOPO™ cloning kit (Invitrogen, USA) and the ligation reaction was allowed to proceed for five minutes are room temperature. Ligated plasmids were transformed into DH5α cells and selected on LB plates supplemented with kanamycin. Colonies were screened for those containing the correct insert via NGS plasmid sequencing. 50% glycerol stocks of final constructs were stored in -80°C.

### Preparation of DNA and RNA reagents for assays

For all RNA reagents other than mat-rRNA, plasmids containing the desired RNA transcript were linearized via incubation with 5% v/v restriction enzymes (MALAT1 – SalI, GALT_1k – NotI / SP6, GALT_full – KpnI, Acc65I / T7, CD44_4k – NotI / SP6, CD44_7k – BamHI / T7) and 10% v/v CutSmart Buffer (New England Biolabs, USA) at 37°C for 4 hours. Linearization was confirmed by gel electrophoresis of DNA samples in a 1% agarose gel with non-digested DNA as controls. Linearized plasmids were purified via PCR clean-up kit (IBI Scientific, USA) per manufacturer manual. In vitro transcription was performed for each transcript using the mMESSAGE mMACHINE Transcription Kit (ThermoFisher, USA) with either SP6 or T7 polymerase as indicated. Transcribed RNAs were purified using the Monarch RNA clean up kit (New England Biolabs, USA). The purity and molecular weight of RNA were confirmed using gel electrophoresis. Confirmed RNA transcripts were aliquoted into single use volumes (2 µL), flash frozen with liquid N2, and stored at -80°C. Mature rRNA (mat-rRNA) is a mixture of rRNAs (18S and 28S), which was purified and labelled with AlexaFluor 647 hydrazide as described (King et al. Cell 2024). MALAT1, CD44, CD44_7k, GALT, and GALT_full transcripts were labelled with AlexaFluor 647 or 488 hydrazide as described (King et al. Cell 2024).

### Protein expression and purification

SRSF1 was expressed in BL21-CodonPlus(DE3)-RIPL E. coli cells (New England Biolabs, USA) grown in LB broth (Millipore Sigma, USA), supplemented with kanamycin, spectinomycin, and chloramphenicol. Cultures were incubated at 37°C in a shaker incubator at 220 rpm until OD600 of 0.4 was reached. Cultures were then induced with 0.35 mM Isopropyl b-D-1-thiogalactopyranoside (IPTG) and incubated for additional 6-8 hours at 23°C. Cells were harvested by centrifugation at 4000 rpm for 30 minutes, washed of in Lysis Buffer (10% glycerol, 20 mM HEPES (pH 8.0), 1.25 M NaCl), and stored as pellets at -80°C.

For lysis, the cell pellet was gently resuspended in 5 mL of cold Lysis Buffer per gram of pellet with one protease inhibitor tablet (Millipore Sigma). 1 mg/mL of lysozyme was added and incubated at 4°C for 1 hour with gentle rocking. The solution was then sonicated (Branson 550 with an L102C horn attachment) using five sets of the following 20-round cycle: 1 second on / 2 second off at 30% power. The solution was then centrifugated for 30 minutes at 10,000 rpm.

Supernatant was incubated with equilibrated 4 mL HisPur Ni-NTA resin (Thermofisher Scientific, USA) at 4°C for 1 hour. Then, the solution was poured into a manual column, and the flow-through fractions were collected. The resin was then washed with 30 mL lysis buffer and the eluted by Elution buffer (10% glycerol, 20 mM HEPES (pH 8.0), 1.25 M NaCl, 300 mM Imidazole) in 2 mL fractions. The fractions were confirmed by SDS-PAGE. Fractions containing His-SUMO-SRSF1 were pooled and cleaved of SUMO tags during an overnight dialysis in the presence of 1:50 molar ratio of His-tagged ULP1 protease to His-SUMO-SRSF1 in Cleavage Buffer (10% glycerol, 20 mM HEPES (pH 8.0), 1.25 M NaCl, 1 mM DTT (dithiothreitol)). The dialysate was incubated with 4 mL of Ni-NTA resin equilibrated with lysis buffer. Flow-through was collected, and the resin was eluted with elution buffer in 5 mL fractions. Presence of SRSF1 was confirmed in the flow-through and His-SUMO and His-Ulp1 in the elution fractions. Flow-through fractions were pooled and diluted 25-fold in dilution buffer (10% glycerol, 20 mM HEPES (pH 8.0)) to lower the total [NaCl]. This solution was further purified using a HiTrap Heparin HP 5 mL column (Cytiva, USA) on a continuous gradient with Buffer A (10% glycerol, 20 mM HEPES (pH 8.0), 50 mM NaCl), and Buffer B (10% glycerol, 20 mM HEPES (pH 8.0), 1.25 M NaCl) by FPLC (ÄKTA Pure, Cytiva, USA). Peak fractions were confirmed by SDS-PAGE, pooled, and concentrated in Amicon Ultra 3 MWCO (molecular weight cut-off) concentrator columns (Millipore Sigma, USA). Concentrated protein was aliquoted into 100 µL, flash frozen in liquid N2, and stored at -80°C. Concentration of protein was determined by absorbance at 280 nm. Predicted extinction coefficient for SRSF1 is 27850 M-1cm-1 (cite Philo, Eur Biophys J 2023).

### Tagging proteins with fluorescent reporters

For covalent conjugation of SRSF1 with NHS(h-hydroxysuccinimide ester) AlexaFluor 488 (Thermofisher Scientific, USA), 1 mg of SRSF1 was dialyzed into Labelling Buffer (10% glycerol, 20 mM HEPES (pH 8.3), 1.25 M NaCl) using Pierce Microdialysis Plates (ThermoFisher Scientific, USA). The dialysate was combined with NHS-Alexa Fluor dye (maintained in DMSO) at a dye:protein molar ratio of 4:1. This mixture was incubated under gentle rocking for one hour at 4°C in the absence of light. The mixture was then dialyzed extensively into Buffer H (10% (v/v) glycerol, 20 mM HEPES (pH 7.4), 50 mM KCl, 5 mM MgCl2) to remove unincorporated dye. Concentrations of protein and AlexaFluor 488 were determined by absorbance measurements (ε280 = 27850 M-1cm-1, ε495 = 73,000 M-1cm-1). Labeling efficiency was determined as the molar ratio of dye to protein and ranged between 0.4 – 0.6. Labeled SRSF1 was aliquoted into single-use volumes (2 µL), flash frozen in liquid N2, and stored at -80 C.

### Confocal microscopy of SRSF1 condensates

Confocal microscopy imaging performed on a Eclipse Ti2 microscope (Nikon, Japa) with a Yokogawa CSU X1 disk module and a LunF laser launch equipped with 405 nm, 488 nm, 561 nm, and 647 nm lasers, using a 60X, 1.4 NA Apo oil immersion objective (Nikon, Japan) and an Orca Flash 4.0 CMOS camera (Hamamatsu, Japan). All images were captured at room temperature using NIS-Elements software (Nikon, Japan) and saved as 16-bit ‘.nd2’ files. Images within a data set were taken with identical imaging parameters ensuring that signal was not saturated (averaging, binning, and projecting were not used). All images of purified SRSF1 and RNA constructs shown are representative crops of one or a few entities (e.g., a condensate) where the brightness and contrast have been optimized. Line scans were obtained using ImageJ and plotted using Prism 10 (GraphPad, USA). Samples were prepared with 1:250 molar ratio of labelled-to-unlabelled SRSF1 and 1:50 labelled-to-unlablled RNA and imaged in silicone wells mounted on a coverslip.

### 90° Light scattering assay

SRSF1 was dialyzed against Buffer H (10% glycerol, 20 mM HEPES (pH 7.4), 50 mM KCl, 5 mM MgCl2) using Pierce Microdialysis Plates (ThermoFisher Scientific, USA) for each set of experiments. The intensity of static right-angle light scattering at 320 nm (I) was measured at each titration point using PTI Quantamaster (Horiba Scientific, Japan). The intersection of linear fits to I versus [SRSF1] (log-log plot) identified discontinuity points in the concentration dependence of the scattering intensity, which is indicative of an assembly formation and/or phase transition. At least two biological and three technical replicates were performed for each experiment.

Optimal linear fits were determined using a modified jackknife approach as described^16^. In summary, each dataset was subjected to two independent series of linear fits, with one series of fits for the low concentration arm (LCA) and the second series of fits for the high concentration arm (HCA). For each arm, a series of fits was initiated with a linear fit to the four lowest or highest concentration data points, respectively, and the root mean square error (RMSE) of each fit was recorded. Following these initial fits, the next highest or lowest concentration data point was added to the respective LCA or HCA data set, the expanded data sets were re-fit, and the new RMSEs were recorded. This process was continued, expanding the fitted dataset by one data point at a time, until all the points in the full dataset were included in both the LCA and HCA linear fits. The intersections of each pair of LCA and HCA best fits were recorded and the average intersection point for all best fits, for all trials at a given concentration of SRSF1, were determined. This is the value reported as the threshold concentration for condensate formation.

Polyphasic linkage is a type of polysteric linkage and describes the influence of ligand (MALAT1 or CTD) binding on the phase behavior of the macromolecule (SRSF1) and is described as follows:

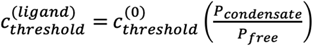

where c(ligand)threshold is the threshold concentration of SRSF1 in the presence of the ligand, c(0)threshold is the threshold concentration of SRSF1 in the absence of ligand, Pcondensate is the binding polynomial of SRSF1 forming condensates, and Pfree is the partition function of free SRSF1 molecules that are not part of the condensates. Each partition function is a sum of macromolecule concentrations either as the condensate or free species referenced to the total monomer concentration of the macromolecule.

## Notes

### Competing Interest Statement

The authors have declared no competing interest.

