## supplementary data for "Chromatin-associated lncRNA-splicing factor condensates regulate hypoxia responsive RNA processing of genes pre-positioned near nuclear speckles"

[1]Department of Cell and Developmental Biology, University of Illinois at Urbana-Champaign, Urbana, IL; [2]Cancer Center at Illinois, University of Illinois at Urbana-Champaign, IL; [3]Department of Biomedical Engineering and Center for Biomolecular Condensates, Washington University in St. Louis, St. Louis, MO; [4]Department of Biochemistry, University of Illinois at Urbana-Champaign, Urbana, IL; [5]Department of Molecular and Integrative Physiology, University of Illinois at Urbana-Champaign, Urbana, IL; [6]Ionis Pharmaceuticals Inc., Carlsbad, CA; [7]Department of Bioengineering, University of Illinois at Urbana-Champaign, Urbana, IL; [8]Carl R. Woese Institute for Genomic Biology- Anticancer Discovery from Pets to People, University of Illinois at Urbana-Champaign, Urbana, IL; [9]Beckman Institute for Advanced Science and Technology, University of Illinois at Urbana-Champaign, Urbana, IL; [10]Division of Biology and Biological Engineering, Caltech, Pasadena, CA.

#: equal contribution

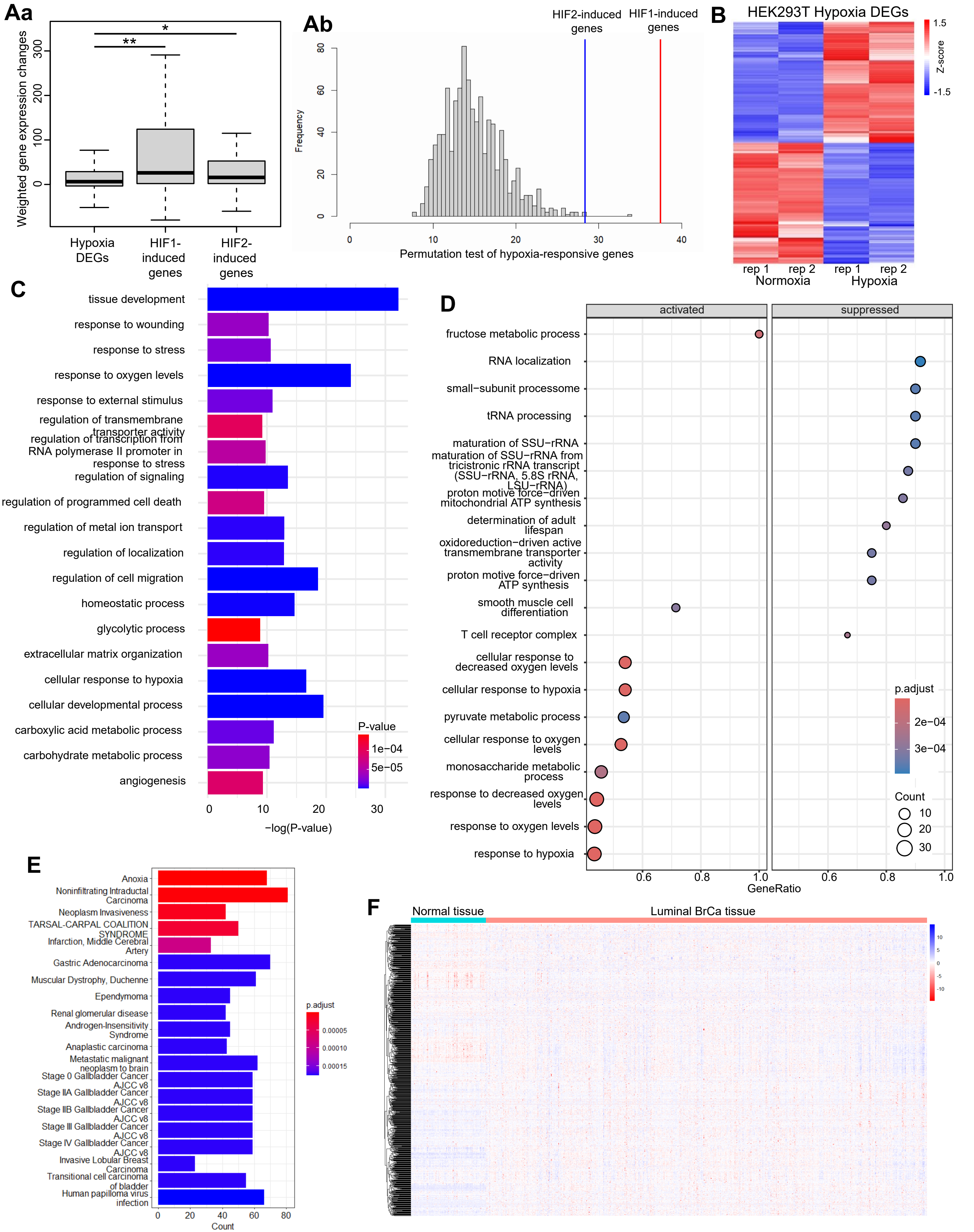


Supplementary Figure 1

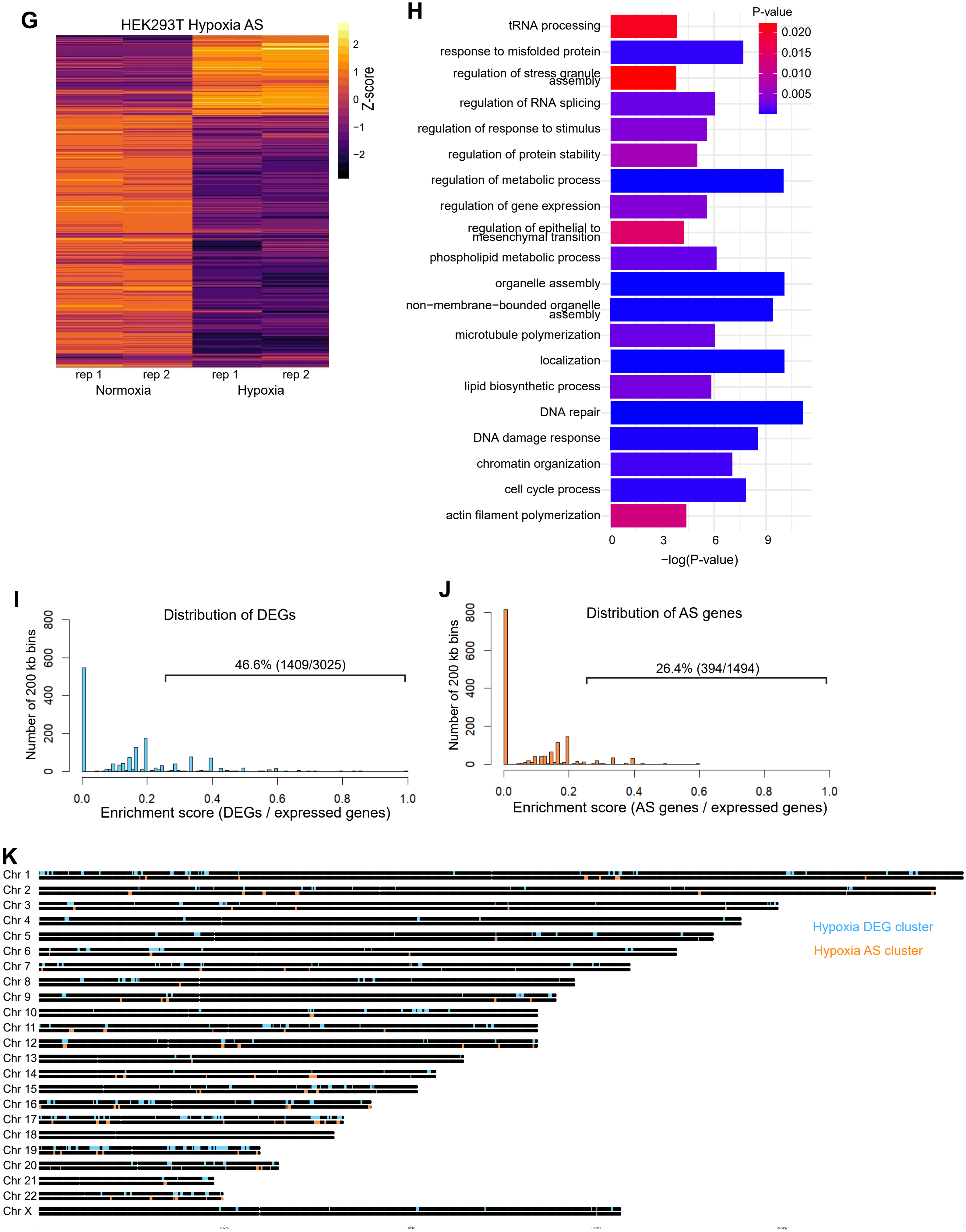


Supplementary Figure 1

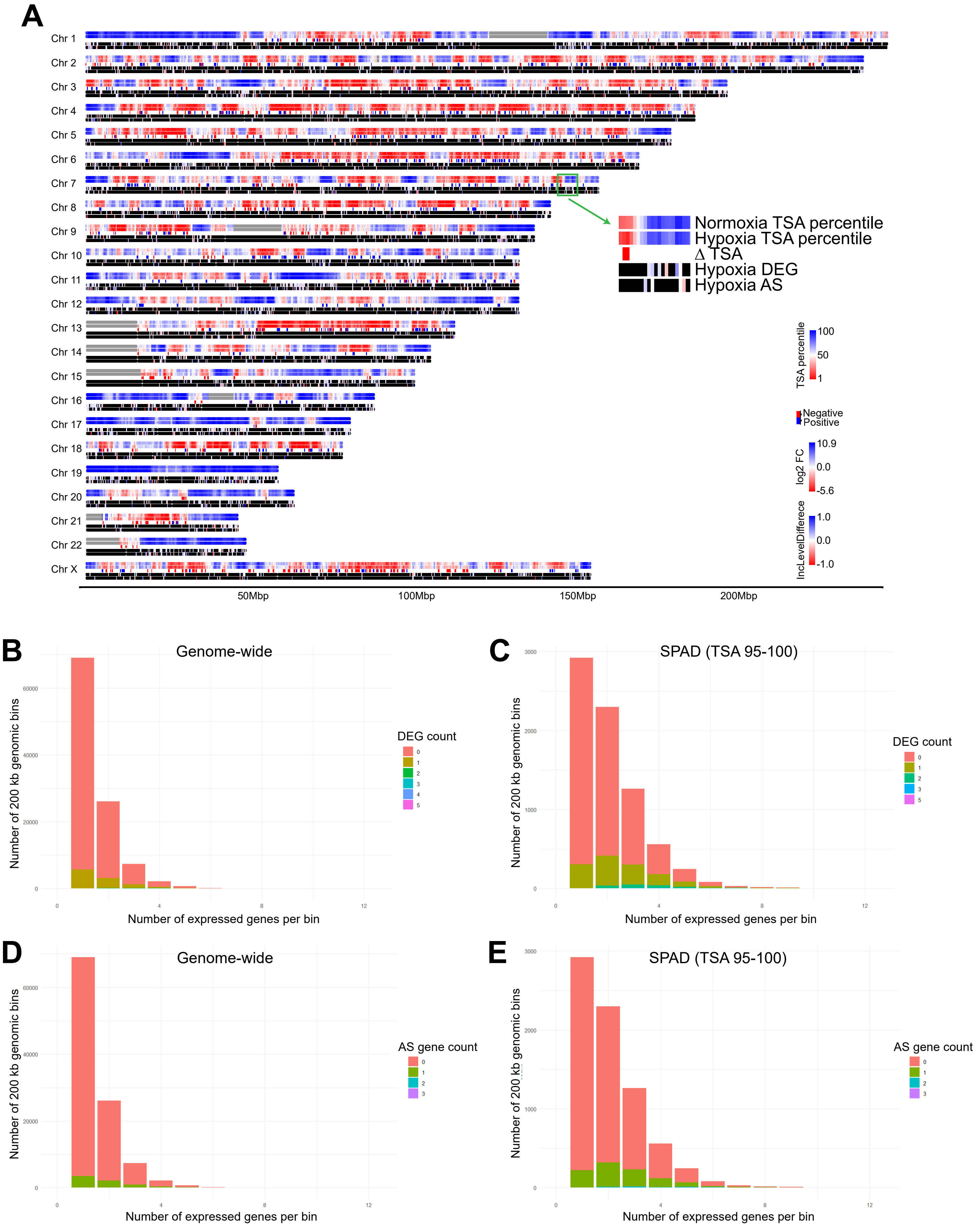


Supplementary Figure 2

F
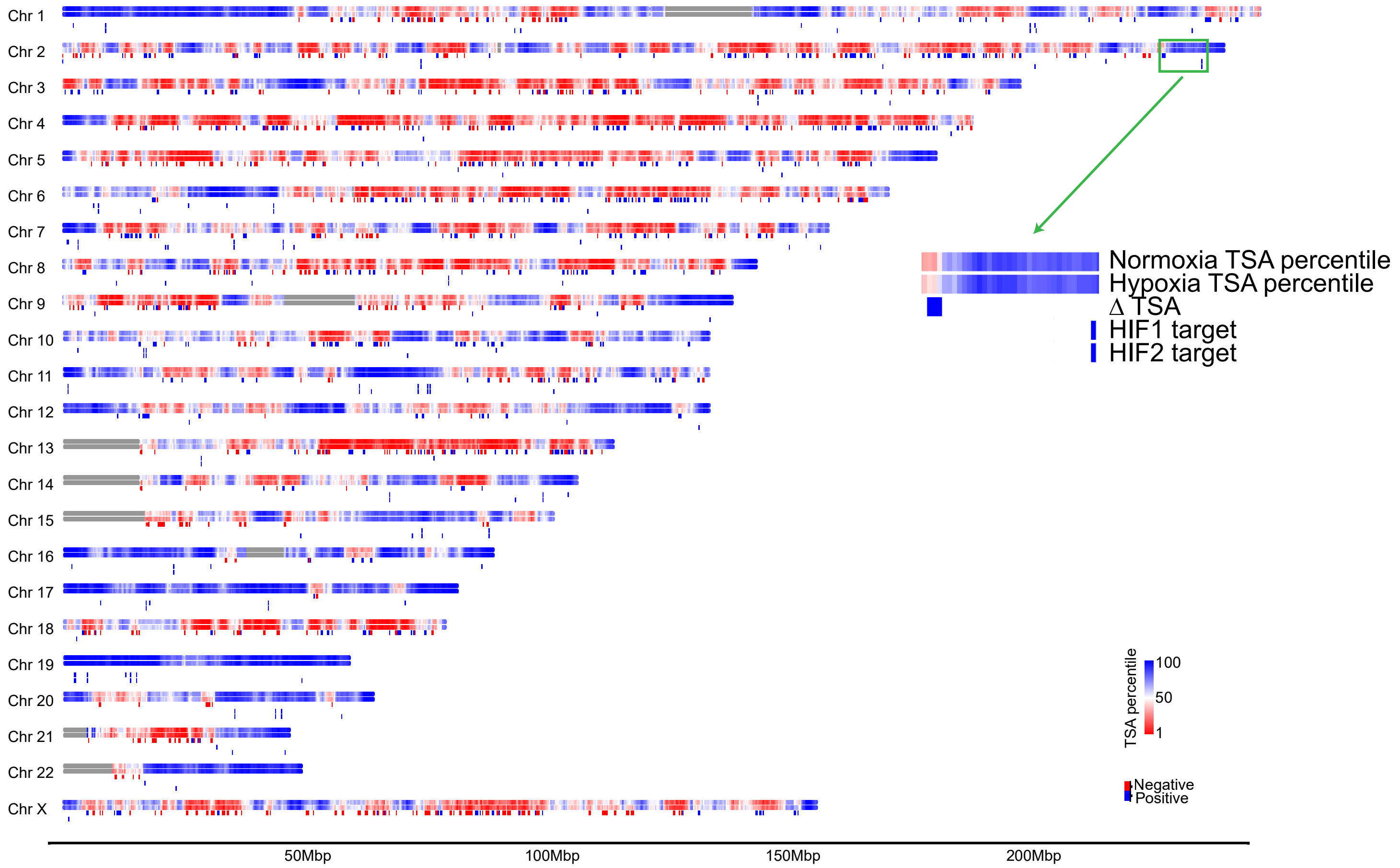


G
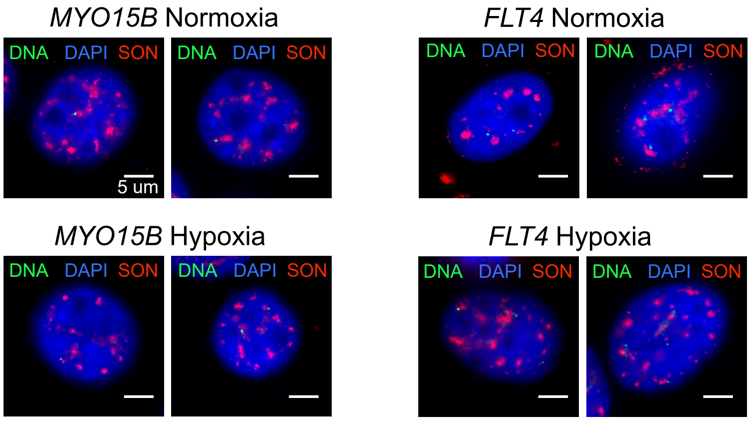


H
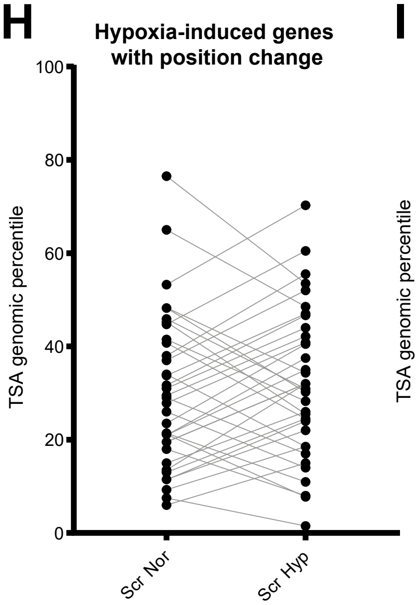


I
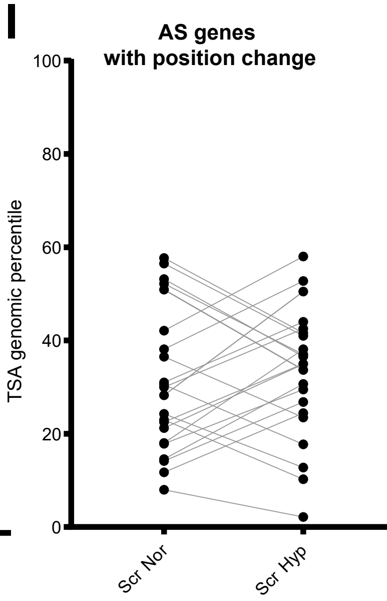


J
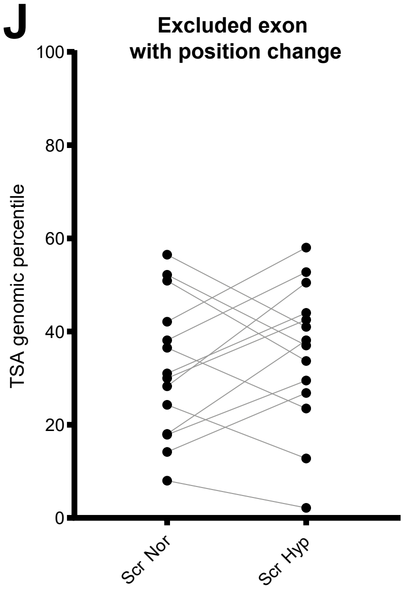


K
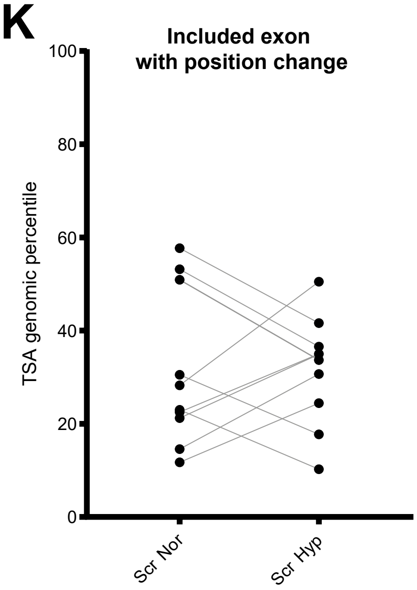


Supplementary Figure 2

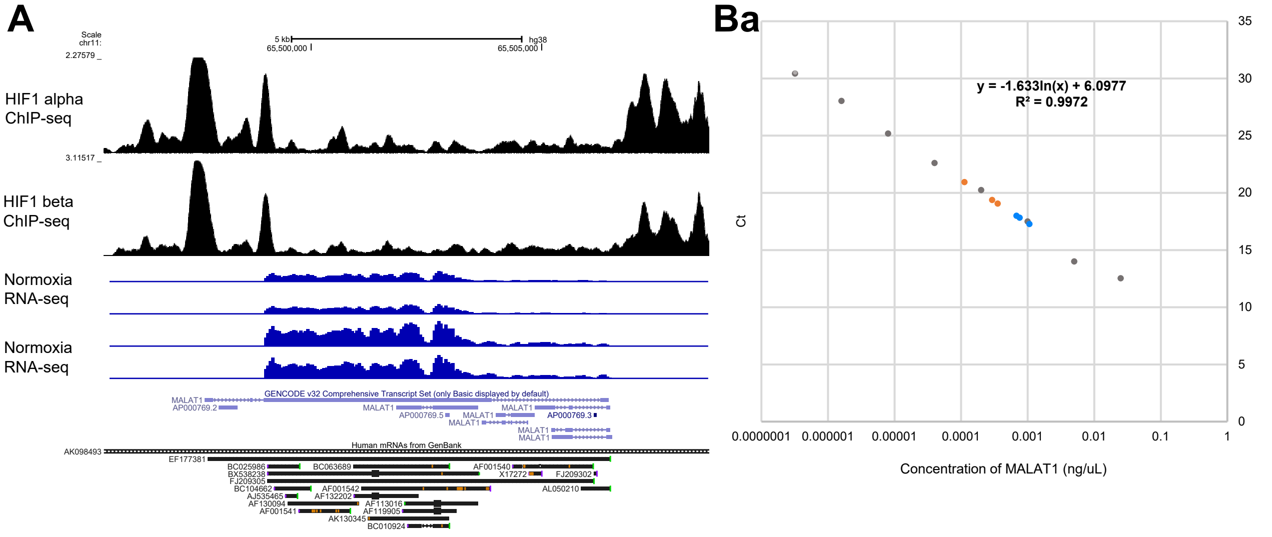


## Bb

| Sample | Ct | Conc. of MALAT1 ng/uL | molecules MALAT1 per cell |
|---|---|---|---|
| Nor-1 | 19.38703 | 0.000292225 | 5492.49933 |
| Nor-2 | 19.06651 | 0.000355599 | 6683.655792 |
| Nor-3 | 20.94901 | 0.000112282 | 2110.390868 |
| Hyp-1 | 18.00782 | 0.000680012 | 12781.14358 |
| Hyp-2 | 17.27636 | 0.001064264 | 20003.33174 |
| Hyp-3 | 17.82893 | 0.00075874 | 14260.86512 |

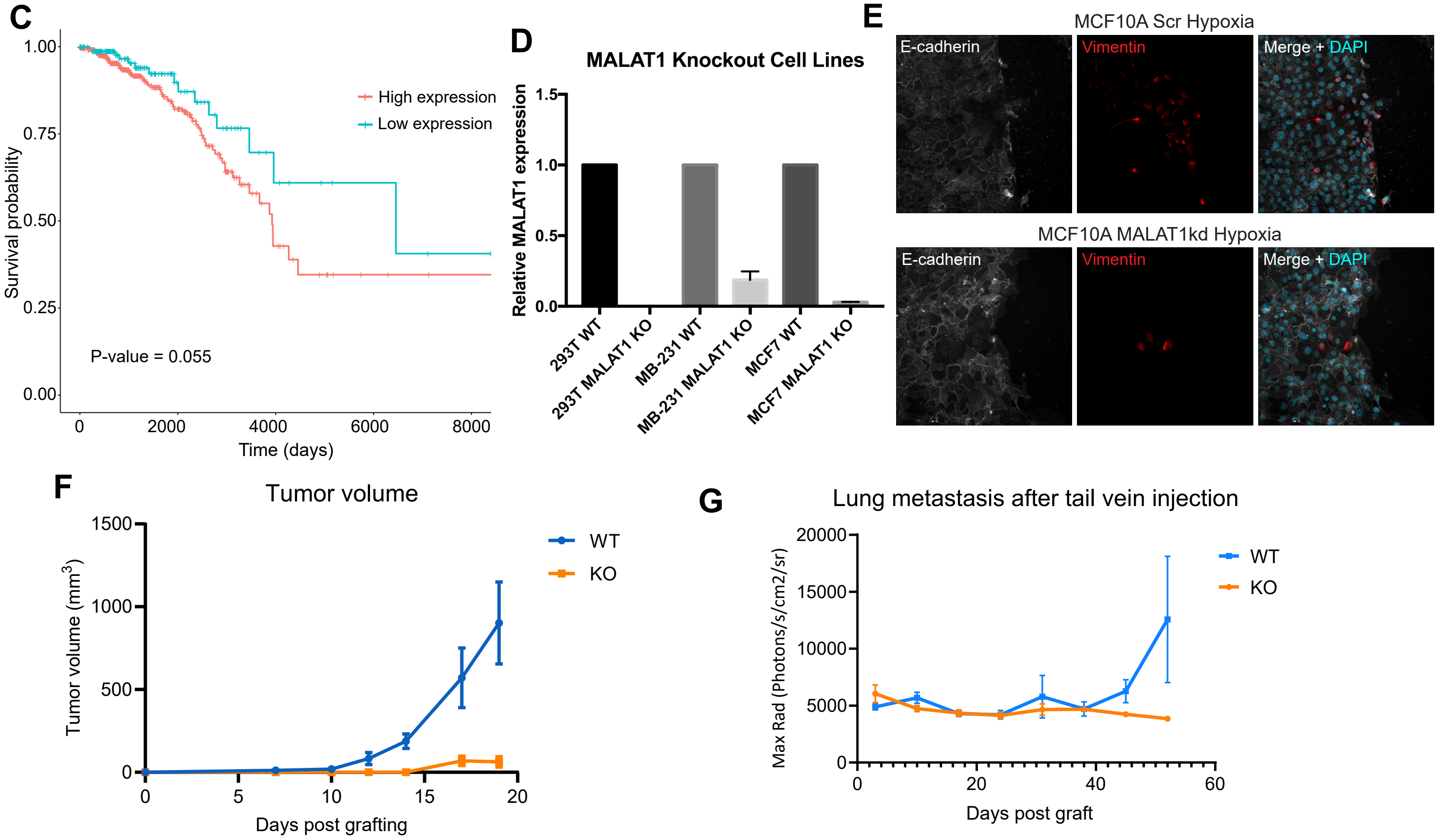


Supplementary Figure 3

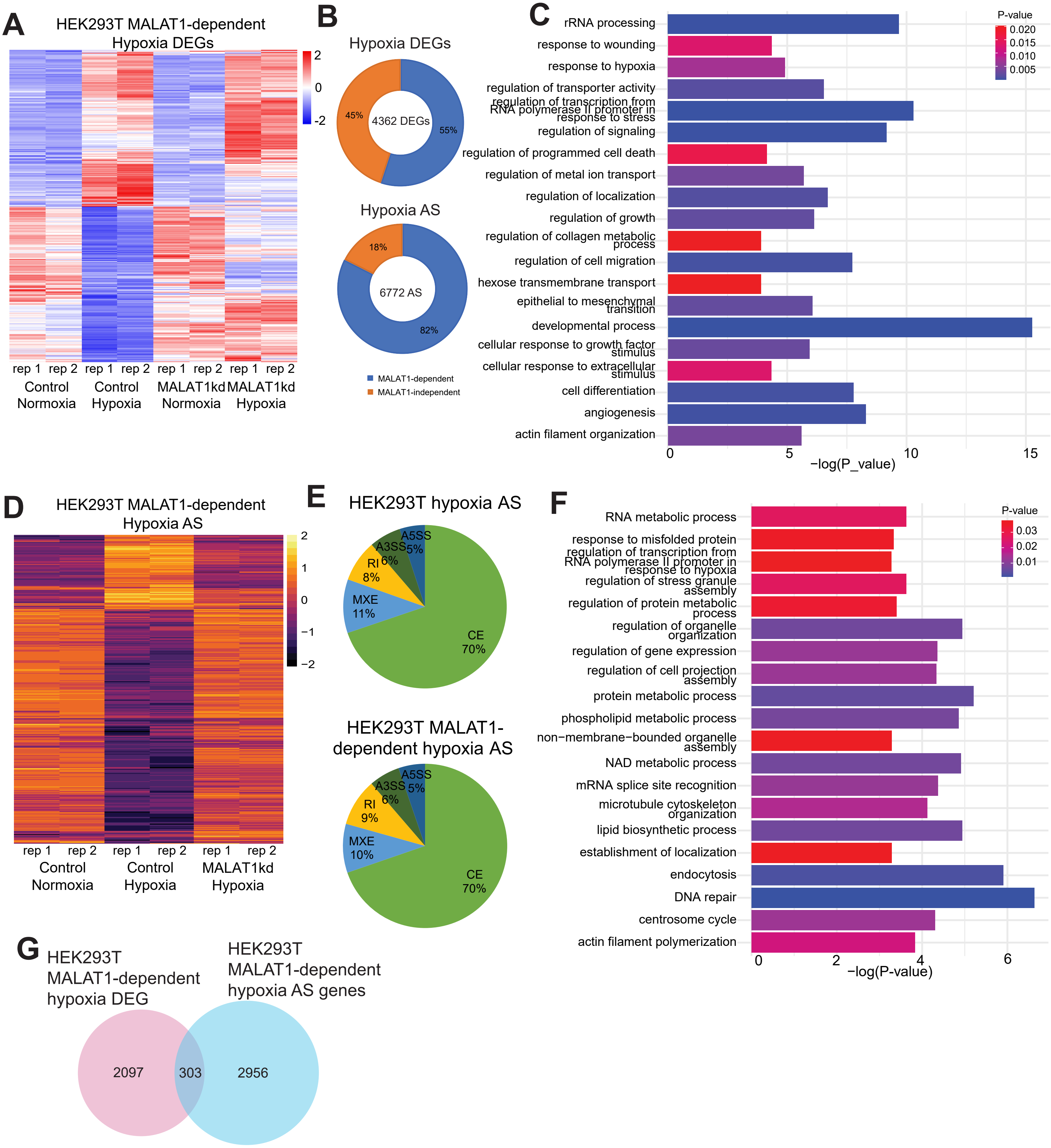


Supplementary Figure 4

H

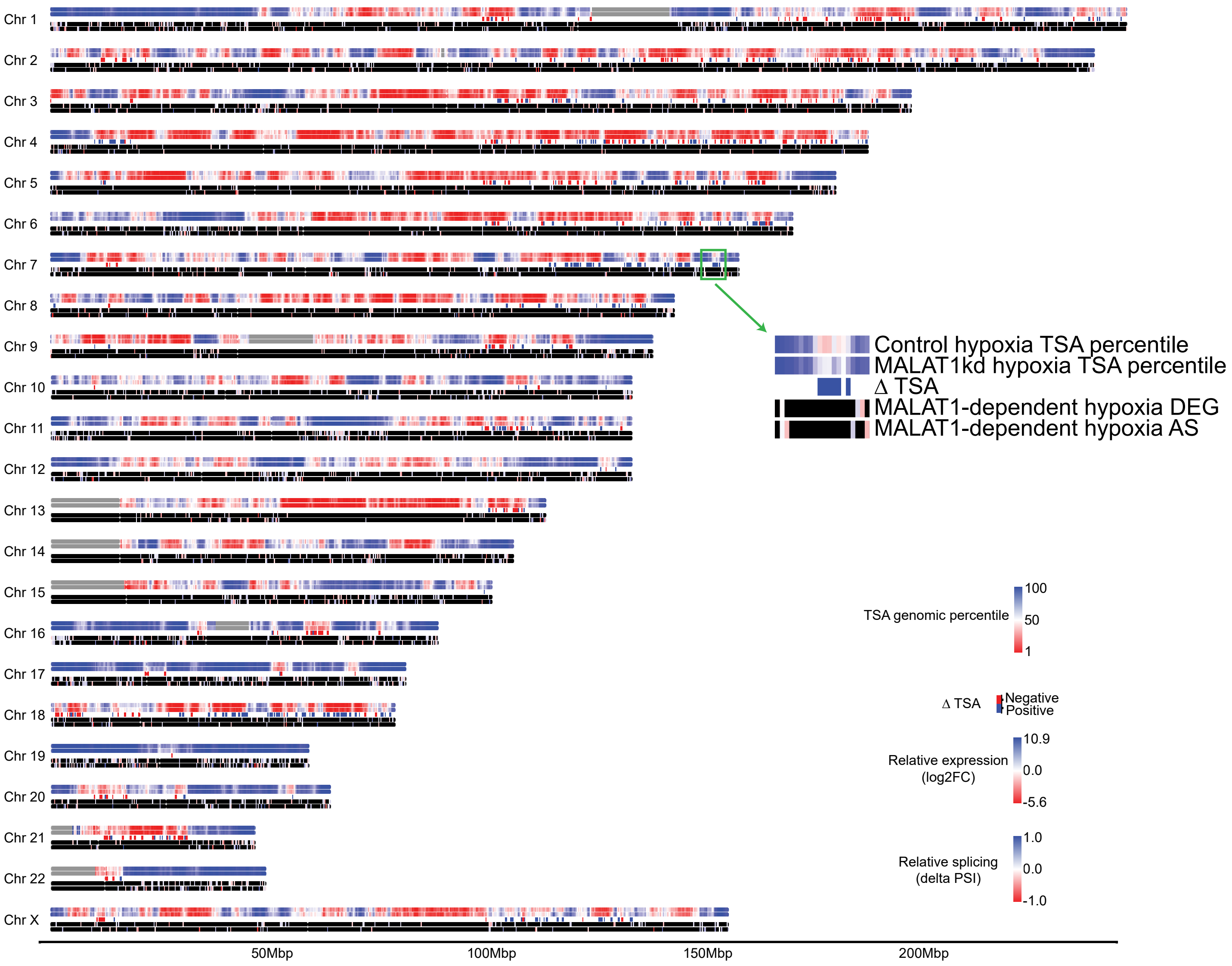


I

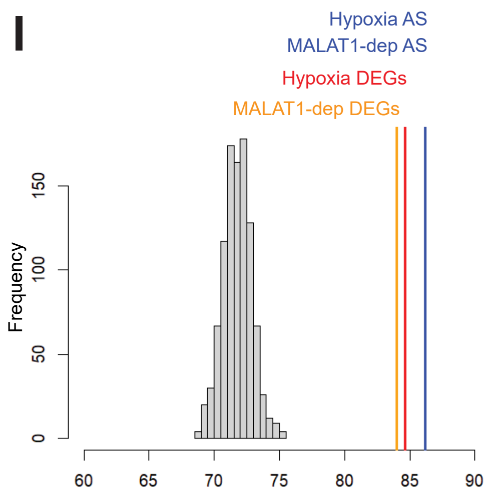


Supplementary Figure 4

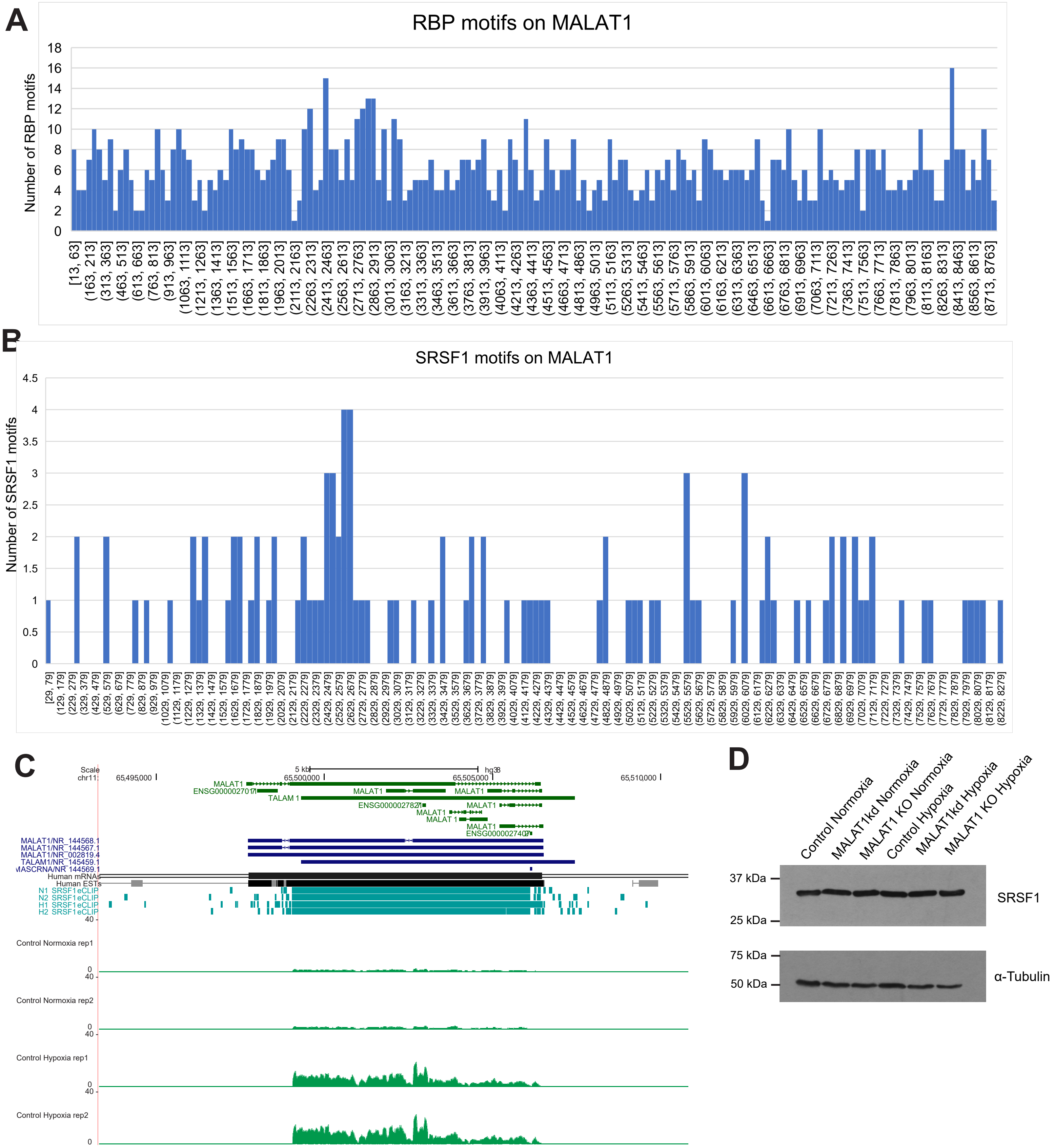


Supplementary Figure 5

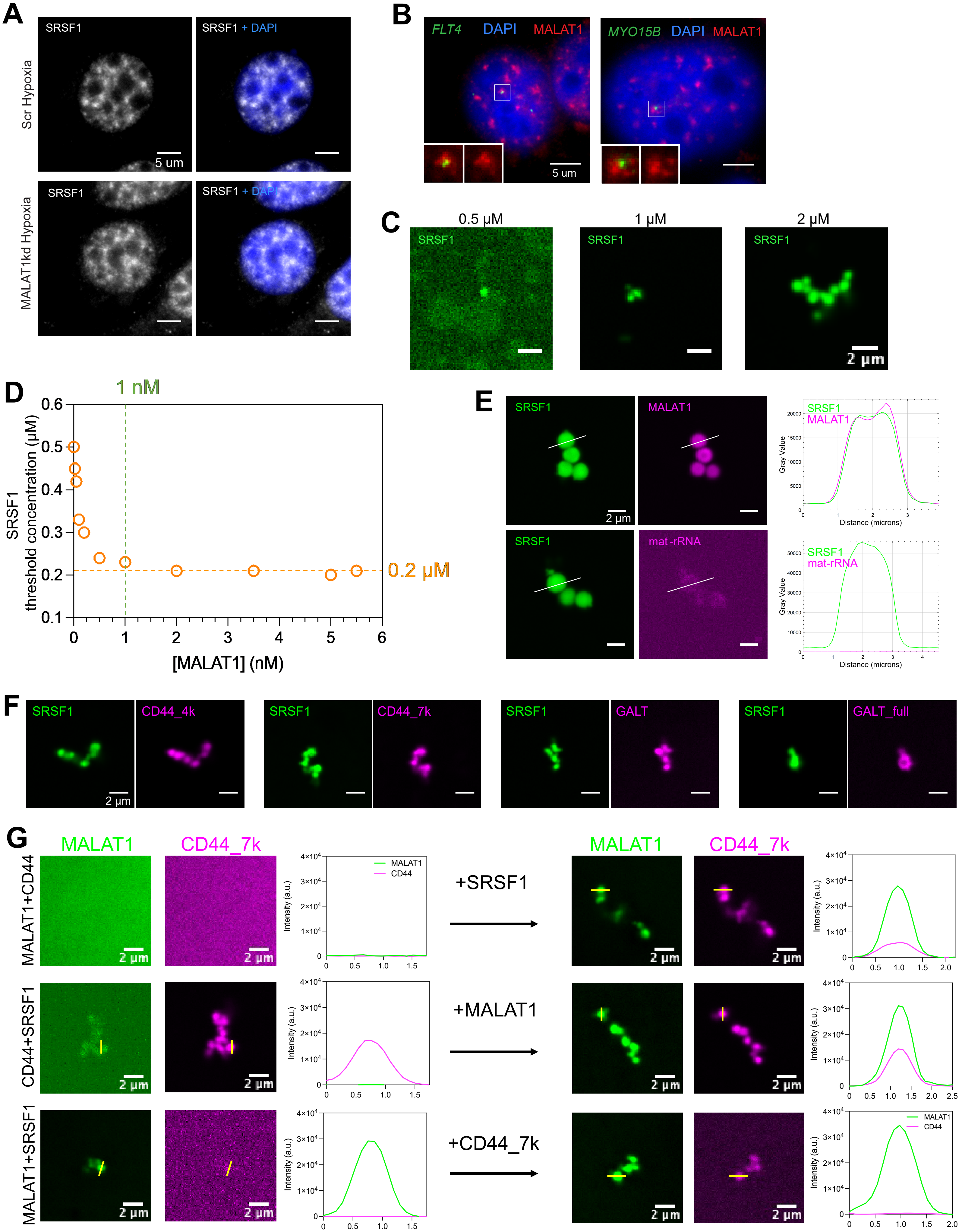


Supplementary Figure 6

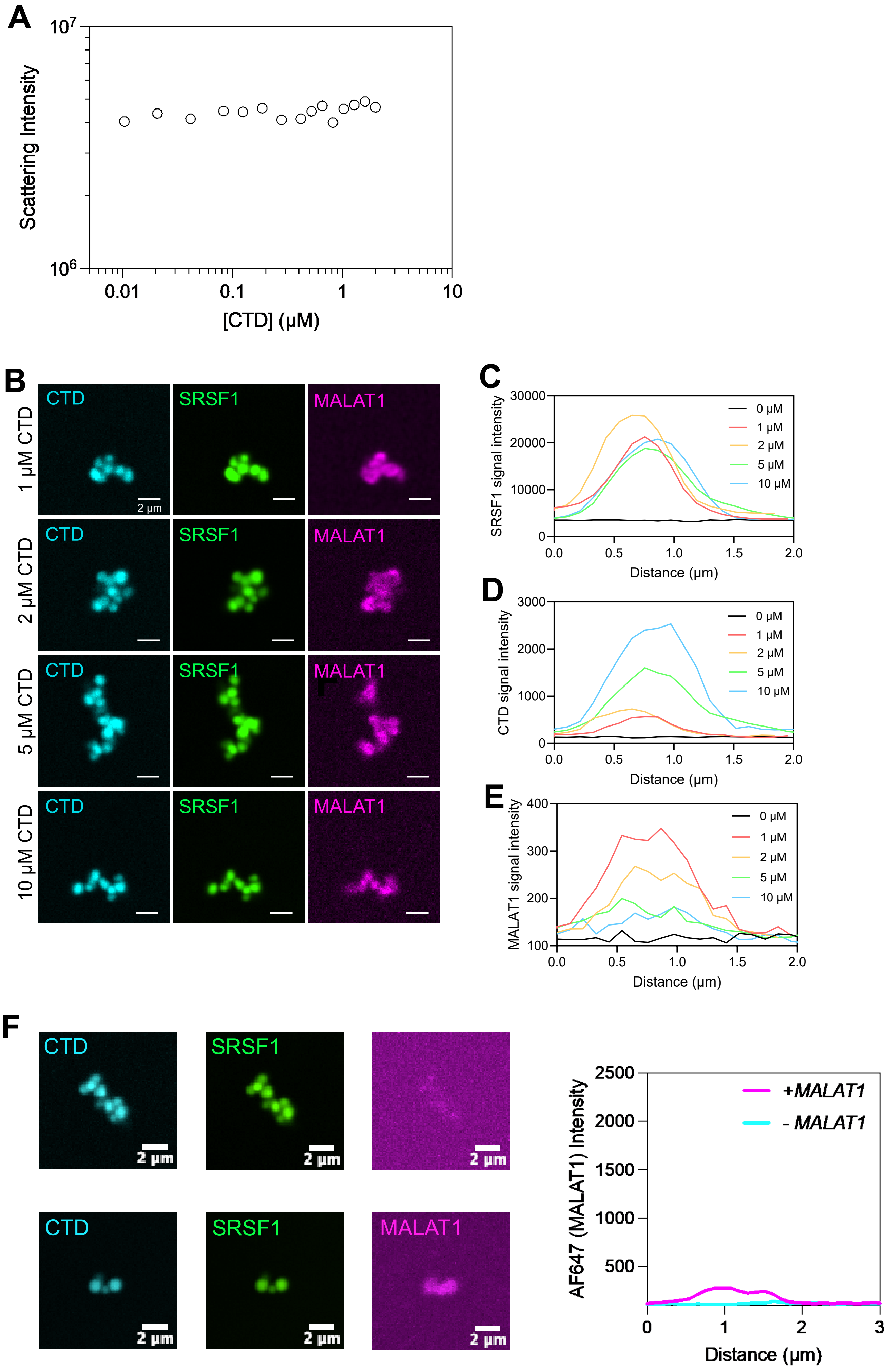


Supplementary Figure 7

Supplementary Figure 1. Hypoxia induces differential gene expression and alternative splicing of pre-mRNA in BrCa cells.

(Aa) Box plot of weighted gene expression changes of HIF1- and HIF2-induced genes in hypoxic MCF7. Box representing 25$^{th}$ to 75$^{th}$ percentile with line indicating mean value. Whiskers representing the min and max value. (Ab) Histogram showing permutation test of weighted gene expression changes of HIF1 and HIF2 target genes compared to all expressed genes in hypoxic MCF7. (B) Heatmap showing the hypoxia-responsive differentially expressed genes in HEK293T cells. Genes (rows of heatmap) are hierarchically clustered using average-linkage clustering method. Color represented in Z-scores are computed per gene by subtracting the mean and then dividing by the standard deviation. Red and blue indicate up-regulated and down-regulated genes respectively. (C) Gene ontology analysis of hypoxia differentially expressed genes in MCF7. (D) Gene set enrichment analysis of hypoxia differentially expressed genes in MCF7. (E) Disease gene set enrichment analysis of hypoxia differentially expressed genes in MCF7. (F) Heatmap showing hypoxia differentially expressed lncRNAs in normal breast tissue (n = 113) and luminal breast cancer tissue (n = 665). Genes (rows of heatmap) are hierarchically clustered using average-linkage clustering method. Color represented Z-scores are computed per gene by subtracting the mean and then dividing by the standard deviation. Red and blue indicate up-regulated and down-regulated lncRNA in tissue samples. (G) Heatmap showing the hypoxia-responsive AS in HEK293T. Splicing events (rows of heatmap) are hierarchically clustered using average-linkage clustering method. Color represented Z-scores are computed per splicing event by subtracting the mean and then dividing by the standard deviation. Yellow indicates exon inclusion and violet indicates exon exclusion. (H) Gene ontology analysis of genes, showing hypoxia-responsive AS in MCF7. (I) Distribution of DEGs in 200 kb genomic bins. Enrichment is calculated by dividing the number of hypoxia DEGs by the number of expressed annotated genes per 200 kb genomic bin. (J) Distribution of hypoxia-responsive AS genes in 200 kb genomic bins. Enrichment is calculated by dividing the number of hypoxia AS genes by the number of expressed annotated genes per 200 kb genomic bin. (K) Genome-wide distribution of hypoxia-responsive DEG (cyan) and AS (orange) gene clusters in MCF7.

Supplementary Figure 2. Hypoxia-responsive genes pre-position near nuclear speckle (NS) neighborhood in MCF7.

(A) Genome-wide heatmap of SON-TSA percentile in normoxia and hypoxia, changes in gene positioning relative to NSs in hypoxia ($\Delta$TSA), hypoxia-responsive gene expression changes, and hypoxia-responsive AS changes in MCF7. (B) Distribution of hypoxia-responsive DEGs in the genome based on number of genes per 200 kb bin. Y-axis indicates number of bins and x-axis indicates number of annotated genes per bin. Bins with no annotated gene are not shown, thus the first bar represents bins with 1 annotated gene. Bar color indicates number of hypoxia DEGs per bin (i.e. first bar showing bins with 1 annotated gene comprise of ~92.5% with no hypoxia DEG and rest of the ~7.5% where the annotated gene is a hypoxia DEG). (C) Distribution of hypoxia-responsive DEGs in speckle-associated domains (SPADs: TSA scores 95-100) based on number of genes per 200 kb bin. Y-axis indicates number of bins and x-axis indicates number of annotated genes per bin. Bins with no annotated gene are not shown. (D)

Distribution of hypoxia-responsive AS genes in the genome based on number of genes per 200 kb bin. Y-axis indicates number of bins and x-axis indicates number of annotated genes per bin. Bins with no annotated gene are not shown. (E) Distribution of hypoxia-responsive AS genes in speckle-associated domains (SPADs: TSA scores 95-100) based on number of genes per 200 kb bin. Y-axis indicates number of bins and x-axis indicates number of annotated genes per bin. Bins with no annotated gene are not shown. (F) Genome-wide heatmap of TSA percentile in normoxia and hypoxia, changes in gene positioning relative to NSs ($\Delta$TSA), and genomic positions of HIF1 and HIF2 target genes. (G) Representative images of DNA FISH and SON immunostaining of hypoxia-responsive genes in normoxic and hypoxic conditions. (H) Paired line plot of hypoxia-induced genes with significant changes in gene position during hypoxia. (I) Paired line plot of hypoxia AS genes with significant changes in gene position during hypoxia. (J) Paired line plot of hypoxia-induced exon exclusion genes with significant changes in gene position during hypoxia. (K) Paired line plot of hypoxia-induced exon inclusion genes with significant changes in gene position during hypoxia.

Supplementary Figure 3. MALAT1, induced during hypoxia, regulates hypoxia-responsive changes in cancer cells.
(A) HIF1 alpha and HIF1 beta ChIP-seq enrichment on MALAT1 gene locus in hypoxic MCF7. IGV representation, showing RNA-seq signal at the MALAT1 locus in normoxic and hypoxic MCF7. (B) Ba) Linear regression graph for absolute quantification of MALAT1 RNA molecules per cell based on qPCR. Bb) Calculation of MALAT1 RNA quantification in normoxic and hypoxic MCF7 cells in biological triplicates. (C) Kaplan-Meier plot of MALAT1 high vs low expressing luminal breast cancer patients in TCGA. (D) Relative MALAT1 levels in MALAT1 KO cells based on RT-qPCR. (E) E-cadherin and vimentin immunostaining in MCF10A control and MALAT1-depleted hypoxic cells after scratch-wound healing assay. DNA is counterstained with DAPI (blue in merge). (F) Tumor volume of flank xenograft of HEK293T wildtype and MALAT1 KO. (G) Luciferase activity measurement of lung metastasis after tail vein injection of MDA-MB-231-luc wildtype and MALAT1 KO.

Supplementary Figure 4. MALAT1 regulates hypoxia-responsive differential expression and AS in MCF7.
(A) Heatmap of hypoxia-responsive DEGs in control and MALAT1kd HEK293T cells. Genes (rows of heatmap) are hierarchically clustered using average-linkage clustering method. Color represented Z-scores are computed per gene by subtracting the mean and then dividing by the standard deviation. Red indicates up-regulated and blue indicates down-regulated genes in hypoxia. (B) Pie charts indicating percentage of hypoxia-responsive DEGs and DS events dependent on MALAT1 in HEK293T. (C) GO analysis of MALAT1-dependent DEGs in MCF7. (D) Heatmap of hypoxia-responsive AS events in control and MALAT1kd HEK293T cells. Splicing events (rows of heatmap) are hierarchically clustered using average-linkage clustering method. Color represented Z-scores are computed per splicing event by subtracting the mean and then dividing by the standard deviation. Yellow indicates exon inclusion and violet indicates exon exclusion. (E) Pie chart indicating classifications of the modes of hypoxia-responsive in HEK293T cells. (F) GO analysis of MALAT1-dependent AS genes in MCF7. (G) Venn

diagram showing the overlap of the MALAT1-dependent hypoxia-responsive DEG and AS genes in HEK293T cells. (H) Genome-wide heatmap of TSA percentile in control hypoxia, MALAT1-depleted hypoxia, changes in gene positioning relative to nuclear speckles in MALAT1-depleted hypoxia ($\Delta$TSA), MALAT1-dependent hypoxia-responsive gene expression changes, and MALAT1-dependent hypoxia-responsive splicing changes. (I) Parametric test of TSA percentiles of hypoxia-responsive and MALAT1-dependent DEG and AS genes compared to overall genome.

Supplementary Figure 5. MALAT1 regulates the SRSF1 binding to its target transcripts. (A) Histogram showing the distribution of RBP motifs across the length of MALAT1 transcript. X-axis represents nucleotide position within MALAT1 where the RBP Motif is present, and Y-axis represents the number of RBP motif at the particular nt position. (B) Histogram showing the distribution of SRSF1 binding motifs across the length of MALAT1 transcript. X-axis represents nucleotide position within MALAT1 where the SRSF1 Motif is present, and Y-axis represents the number of SRSF1 binding motifs at the particular nt position. (C) Genome browser view showing SRSF1 eCLIP peaks on the MALAT1 locus and increase MALAT1 RNA levels in hypoxic HEK293T cells. (D) Immunoblot showing SRSF1 levels in control, MALAT1 kd, and MALAT1 CRISPR KO in normoxic and hypoxic MCF7 cells. $\alpha$-Tubulin is used as loading control.

Supplementary Figure 6. MALAT1 promotes SRSF1 condensate assembly.
(A) Representative image of SRSF1 immunostaining in control and MALAT1kd hypoxic MCF7 cells. DNA is counterstained with DAPI (blue). (B) Representative images of MALAT1 RNA FISH (red) and DNA FISH (green) of MALAT1-regulated genes in hypoxic MCF7. DNA is counterstained with DAPI (blue). (C) Confocal images of SRSF1 in vitro condensates formed at different concentrations of SRSF1. (D) Graph showing SRSF1 threshold concentration over range of MALAT1 concentrations by 90 degree light scattering assay. (E) Confocal images and line profiles for SRSF1 condensates with MALAT1 and mature rRNA. (F) Confocal images of SRSF1 condensates with CD44_4kb, CD44_7kb, GALT 1 kb, and GALT full length RNA. (G) Confocal images and line profiles showing the status of SRSF1 condensates after sequential addition of SRSF1 to MALAT1 + CD44_7k (top), MALAT1 to CD44_7k + SRSF1 (middle), and CD44_7k to MALAT1 + SRSF1 (bottom).

Supplementary Figure 7. MALAT1 promotes the RNA pol II-SRSF1 splicing condensate assembly.
(A) 90 degree light scattering assay of CTD. (B) Confocal image and line profile of SRSF1-CTD condensates without or with the addition of MALAT1. (C) Confocal images of SRSF1 condensates incubated with MALAT1 and increasing concentrations of CTD. (D) Line profile showing SRSF1 (D), CTD (E), and MALAT1 (F) signal intensities in the SRSF1 condensates with increasing concentrations of CTD.

Supplementary Tables

Supplementary Table 1:
Hypoxia DEGs in MCF7

Supplementary Table 2:
Hypoxia DEGs in HEK293T

Supplementary Table 3:
DEGs in Luminal BrCa tissue vs adjacent normal tissue

Supplementary Table 4:
Hypoxia-responsive AS in MCF7

Supplementary Table 5:
Hypoxia-responsive AS in HEK293T

Supplementary Table 6:
HIF1 target genes in MCF7

Supplementary Table 7:
HIF2 target genes in MCF7

Supplementary Table 8:
Hypoxia-responsive DEG hubs

Supplementary Table 9:
Hypoxia-responsive AS hubs

Supplementary Table 10:
MCF7 hypoxia-responsive DEG TSA genomic percentiles

Supplementary Table 11:
MCF7 hypoxia-responsive AS TSA genomic percentiles

Supplementary Table 12:
Hypoxia-responsive lncRNAs

Supplementary Table 13:
MCF7 MALAT1-dependent hypoxia-responsive DEGs

Supplementary Table 14:
MCF7 MALAT1-dependent hypoxia-responsive AS

Supplementary Table 15:
HEK293T MALAT1-dependent hypoxia-responsive DEGs

Supplementary Table 16:
HEK293T MALAT1-dependent hypoxia-responsive AS

Supplementary Table 17:
MCF KO MALAT1-dependent hypoxia-responsive AS

Supplementary Table 18:
RBP motifs on MALAT1 transcript

Supplementary Table 19:
RBP motif enrichment on MALAT1-dependent spliced regions

Supplementary Table 20:
RBP motif enrichment on control spliced regions

Supplementary Table 21:
HEK293T SRSF1-dependent hypoxia-responsive DEGs

Supplementary Table 22:
HEK293T SRSF1-dependent hypoxia-responsive AS

Supplementary Table 23:
MCF7 SRSF1 eCLIP peaks in hypoxia

Supplementary Table 24:
HEK293T SRSF1 eCLIP peaks in hypoxia
